# Brain-derived extracellular vesicle microRNAs in Lewy body and Alzheimer’s disease

**DOI:** 10.1101/2025.06.06.656900

**Authors:** Stephanie J. Yang, Andrew A. Lin, Hanfei Shen, Laura W. Pappalardo, Griffin B. Spychalski, Jean Rosario, Leah K. Forsberg, Kiera M. Grant, Rodolfo Savica, Sid O’Bryant, David T. Jones, Dennis W. Dickson, R. Ross Reichard, Aivi T. Nguyen, David F. Meaney, Bradley F. Boeve, Owen A. Ross, Pamela J. McLean, David Issadore

## Abstract

**INTRODUCTION:** Robust plasma-based biomarkers to distinguish Lewy body disease (LBD) and Alzheimer’s disease (AD) are currently lacking. We applied track-etch magnetic nanopore (TENPO) sorting for enrichment of brain-derived extracellular vesicle (EV) signatures as potential biomarkers to address this gap.

**METHODS:** We analyzed plasma from 137 autopsy-confirmed patients [30 LBD, 31 AD, 30 AD/LBD, 19 AD with amygdala Lewy bodies (AD/ALB), and 27 controls], sequencing miRNAs from TENPO-isolated GluR2-positive (neuron-enriched) and GLAST-positive (astrocyte-enriched) EVs, and measuring plasma proteins (Aβ40, Aβ42, tau, p-Tau181, p-Tau231) via SIMOA.

**RESULTS:** We identified 16 GluR2+, 8 GLAST+, and 4 protein biomarkers with differential expression (false discovery rate-corrected *P* value < .1) between LBD and AD. A multimodal 15-feature panel classified LBD versus AD with 10-fold cross-validated accuracy = 0.95 and area under the curve (AUC) = 0.96.

**DISCUSSION:** Brain-derived EVs offer accurate and accessible miRNA biomarkers for the differential diagnosis of LBD and AD.

## 1 BACKGROUND

Differentiating between Lewy body disease (LBD) and Alzheimer’s disease (AD), the two most prevalent histopathologies underlying neurodegenerative dementia,^1^ is a challenging clinical question affecting patient outcomes and hindering therapeutic development. LBD and AD share overlapping clinical manifestations, despite distinct pathophysiological mechanisms.^2^ LBD presents clinically as dementia with Lewy bodies (DLB), Parkinson’s disease (PD), or Parkinson’s disease dementia (PDD), and is characterized by α-synuclein (α-syn) aggregates,^1^ whereas AD is marked by amyloid β (Aβ) plaques.^3^ While recent efforts have yielded promising biomarkers for diagnosing AD,^4–6^ few non-invasive biomarkers to date are capable of accurately diagnosing LBD or differentiating between LBD and AD.^1, 7–9^ Clinical diagnostic accuracies for LBD remain low, ranging from 34-89.5% when compared to associated postmortem pathological diagnoses,^10–13^ a challenge accentuated by the approximately 50% of LBD and AD patients who exhibit mixed pathology at death.^1, 3, 8^

The incorrect classification of neurodegenerative dementia can have significant impacts on patient outcomes, such as adverse side effects in patients with LBD who receive anticholinergic and conventional antipsychotic medications.^2, 11^ Recently-approved AD therapeutics, such as lecanemab and donanemab, are also more effective when administered early in the disease course.^14^ Moreover, improved LBD and AD biomarkers can accelerate and lower the cost of clinical trials by enriching the patient population for those most likely to benefit, which is critical, given the current absence of disease-modifying therapies for LBD.^1^

The current standard of care for differentiating LBD versus AD primarily relies on (1) clinical assessment, which lacks specificity; (2) neuroimaging, which is not always accessible and has limited sensitivity/specificity^8, 15, 16^; or (3) cerebrospinal fluid (CSF) analysis, which requires invasive lumbar punctures.^5, 17, 18^ By contrast, blood-based biomarkers are minimally invasive and more accessible. Numerous plasma biomarkers have shown potential in differentiating neurodegenerative dementias.^6, 19–22^ However, (1) prior studies rely on clinical diagnosis rather than postmortem pathology as the ground truth, limiting their accuracy;^19, 21, 22^ (2) biomarkers that have shown efficacy in diagnosing dementia from controls show poor performance in classifying between different dementias;^8, 9, 23^ and (3) many circulating biomarkers degrade rapidly in blood, and can be non-specifically released from non-neurological sources or comorbidities.^23^

Extracellular vesicles (EVs) offer a promising solution to the limitations of conventional biomarkers for neurodegenerative disease. Specifically, EVs not only carry surface proteins, internal proteins, and nucleic acid cargoes reflecting their cells of origin^24^ but also can cross the blood-brain barrier (BBB),^25^ therefore representing a source for peripherally circulating brain-derived biomarkers. Multiple studies have interrogated EV cargoes for LBD and AD diagnosis;^26^ despite early promise, these studies have been limited by the absence of enrichment for brain-specific EVs from the large background in blood (10^10^-10^12^ EVs/mL),^27^ low cohort sizes (N < 50 total subjects), or a lack of postmortem pathology.^7, 28–30^

We address these challenges by performing a multimodal analysis of brain-derived EV miRNA cargo and conventional AD protein biomarkers (Aβ40, Aβ42, tau, p-Tau181, p-Tau231)^6^ from the antemortem plasma of 137 subjects with postmortem-confirmed LBD, AD, or mixed pathology, and controls. We leverage previous work from our group, track-etch magnetic nanopore (TENPO) sorting,^27^ a microfluidic technology that isolates tissue-specific EV subpopulations based on sufficient surface protein expression. TENPO achieves high specificity and throughput by massively parallelizing magnetic sorting of individual EVs (through 10^7^ nanopores/chip),^27^ with previously demonstrated utility in brain injury.^31–34^ Within the current study, we isolate two subpopulations of EVs from each sample, using GluR2 to enrich for neuron-derived EVs and GLAST to enrich for astrocyte-derived EVs, capturing a more comprehensive state of the brain than possible with a single pulldown. We combine TENPO with next-generation miRNA sequencing and gold-standard digital enzyme-linked immunosorbent assay (ELISA) to: 1) identify differentially expressed biomarkers between disease cohorts across multiple modalities; 2) assess the degree to which each compartment of biomarkers (GluR2+ EVs, GLAST+ EVs, and proteins) yields complementary diagnostic information; and 3) validate the hypothesis that a multimodal biomarker panel can improve dementia diagnosis. This work articulates a proof-of-concept approach toward developing non-invasive, accessible diagnostic tools for differentiating LBD and AD.

## 2 METHODS

### 2.1 Study population and sample collection

Participants were recruited as part of the Mayo Clinic Alzheimer’s Disease Research Center (ADRC) or Mayo Clinic Study of Aging (MCSA) between 2004 and 2019. Subjects selected for this study were chosen based upon postmortem pathology for LBD,^11^ AD,^35, 36^ mixed AD and LBD (AD/LBD), AD with amygdala Lewy bodies (AD/ALB),^37^ or absence of AD and Lewy body pathology (controls).

Blood samples were collected during multiple visits for most patients. The sample selected for this study was (1) defined as early in the course of the disease (preferably within the initial 1-2 years of participation), (2) one for which sufficient plasma was available, and (3) taken from a patient for which autopsy was performed. For controls, samples were selected using the same criteria with preference being within the initial 1-2 years of participation.

Ethylenediamine tetraacetic acid (EDTA) plasma samples were collected following an overnight fast. Samples were centrifuged, and plasma was aliquoted into polypropylene tubes and stored at −80°C until TENPO EV isolation. During the same visit as sample collection, detailed clinical data was also recorded, including demographic information, cognitive assessment measurements, neuroimaging, and clinical diagnoses; see Diaz-Galvan P et al., *Alzheimers Dement.*, 2024 for additional details.^38^

Postmortem brain tissue was collected, processed, and analyzed via standard procedures as described in Lowe VJ et al., *Alzheimers Dement.*, 2024.^39^ Neuropathologic diagnoses were determined as described above.

### 2.2 Isolation of EVs via TENPO

EV isolation was performed using TENPO chips, consisting of three layers of track-etched nickel-iron coated membranes with 3 μm diameter pores, and assembled via a protocol previously specified in our published work.^27^ Briefly, 500 μL of patient plasma was incubated for 20 minutes at a concentration of 1 μg/mL with either biotinylated GluR2 capture antibody (GluR1 + GluR2 polyclonal antibody, Bioss bs-10042R-Biotin) for neuron-derived EVs per our previous work^32^ or biotinylated GLAST capture antibody [GLAST (ACSA-1) antibody, anti-human/mouse/rat Biotin, Miltenyi Biotec, 130-118-984] for astrocyte-derived EVs, where the use of GLAST as a protein target for astrocyte EV isolation has been previously reported.^40, 41^ The sample was then incubated with 50 μL of 50 nm magnetic nanoparticles (MNPs) at manufacturer stock concentration (Anti-Biotin MicroBeads UltraPure, Miltenyi Biotec) for another 20 minutes. During incubation, the TENPO devices were pre-blocked with 700 μL 1% Pluronic F-127 before a 1 mL phosphate-buffered saline (PBS) wash at 15 mL/hr using a syringe pump (Braintree Scientific). Following off-chip sample incubation and device blocking/washing, sample was added on-chip while on magnet and flowed through at 1 mL/hr. 100 μL PBS was added once the entire sample had passed through the chip. The device was then washed three times with 700 μL PBS at 15 mL/hr to remove non-specific background.

### 2.3 RNA isolation from EVs

To isolate RNA from the EVs captured on-chip, 700 μL of Qiazol (Qiagen) was added to the device and incubated for 2 minutes on-chip for lysis and release of RNA cargo. Lysate was then collected at 15 mL/hr and frozen at -80°C until RNA isolation. RNA isolation was conducted using the miRNeasy Mini Kit (Qiagen) per manufacturer instructions; isolated RNA (20 μL elution volume) was aliquoted into three aliquots and frozen at -80°C until sequencing preparation.

### 2.4 Next-generation miRNA sequencing

Isolated RNA was prepared into libraries for miRNA sequencing using the Qiagen QIAseq miRNA library kit (96) following manufacturer instructions. To prepare 274 miRNA samples (two samples for each of the 137 patients - one per EV pulldown), library preparation was conducted in four rounds. Libraries were then barcoded using the QIAseq miRNA 96 Index IL (96). The concentration of each library was assessed using a Qubit fluorometer (dsDNA HS Kit, Invitrogen), and library size distribution for selected libraries was assessed using a Bioanalyzer HS DNA Analysis kit (Agilent). Libraries were sequenced on the Novaseq 6000 at the Next-Generation Sequencing Core at the University of Pennsylvania using the Novaseq S1 kit (100 cycles) (Illumina) per manufacturer instructions. Libraries and 1% PhiX spike-in (Illumina) were pooled based on Qubit readings and diluted to 2 nM in Trizma (Sigma-Aldrich). 100 μL of diluted library alongside 1 μL diluted PhiX were denatured with 25 μL of 0.2 N NaOH before neutralization with 25 μL of 400 mM Tris-HCl and loading into the sequencer for single-read sequencing per Qiagen instructions. For all sequencing runs following the first run, overlapping libraries from previous batches were re-run to check for batch-to-batch variation. Comparison of the libraries run on different batch days showed average Kendall correlation > 0.9 for the sequencing reads from the same patient sample run on two different batch days (**Supplementary Fig. 1**), as well as close rank-order matching of individual patient samples between sequencing runs (**Supplementary Fig. 2**).

### 2.5 RNA sequencing data analysis pipeline

RNA sequencing data was demultiplexed via Illumina BCL Convert before alignment, counting, and counts-per-million (CPM) normalization using the Qiagen RNA-seq Analysis Portal. The Qiagen RNA-seq Analysis Portal was additionally used to conduct tests of differential expression between disease cohorts, using Wald tests based on negative binomial distributions, and multiple hypothesis testing correction, using the Benjamini-Hochberg method with the Qiagen default false discovery rate (FDR) of 0.1. Pathway analysis was conducted via DIANA miRPath v4.0 using the TarBase v8.0 database (accessed September 2024).^42^ Data visualization, calculation of receiver operating characteristic (ROC) and area under the curve (AUC), and evaluation of least absolute shrinkage and selection operator (LASSO) panels using an ensemble model (which included k-nearest neighbors, support vector machine, linear discriminate analysis, logistic regression, and Naive Bayes, based on our prior work)^43^ were conducted using MATLAB R2024a (MathWorks). LASSO biomarker selection was implemented using Python. Delong’s test was applied via a framework adapted from Sun X and Xu W, *IEEE Signal Process. Lett.*, 2014,^44^ for which biomarkers were defined as significant relative to a non-predictive biomarker (AUC = 0.5) at *P* value < .1.

### 2.6 Plasma protein analysis

Analysis of protein in patient plasma was conducted via the Neuro3Plex (Aβ40, Aβ42, tau), p-Tau181, and p-Tau231 Single MOlecule Array (SIMOA) digital ELISA assays from Quanterix. All SIMOA assays were run on an HD-X instrument per manufacturer instructions at the Human Immunology Core at the University of Pennsylvania. Samples were run in triplicate and fitted to a standard curve to yield protein concentrations. For samples outside of the dynamic range, protein concentrations were set to either the minimum fitted concentration on-plate (if below dynamic range) or upper assay range as reported by Quanterix (if above dynamic range). The Aβ42/Aβ40 ratio for each patient was calculated using their Aβ42 and Aβ40 concentrations and incorporated into downstream differential analysis.

### 2.7 Nanoparticle tracking analysis (NTA) of TENPO-isolated EVs

Plasma from eight different patients (62.5 µL per patient) was pooled to create a 500 µL composite patient sample for each of the five disease cohorts investigated in this study. EVs were eluted off-chip using a previously reported protocol.^27^ NTA was performed at the Extracellular Vesicle Core at the University of Pennsylvania using the Particle Metrix ZetaView instrument. All dilutions were performed on-site in deionized water.

### 2.8 Isotype control validation of GluR2 and GLAST antibodies

To confirm the specificity of the GluR2 and GLAST antibodies, quantitative polymerase chain reaction (qPCR) was performed on miRNA from EVs captured by TENPO using either 0.5 µg GluR2 or 0.75 µg of GLAST capture antibody, alongside the appropriate isotype control antibodies at the same respective concentrations (rabbit IgG isotype control antibody, BioLegend, 910801 for GluR2; mouse IgG2a isotype control antibody, Miltenyi, 130-106-546 for GLAST) on 500 µL of pooled healthy human plasma (BioIVT). To demonstrate specificity of the antibodies for EV miRNAs associated with the central nervous system (CNS), we used a commercially available plate containing probes for 84 miRNAs known to be associated with EVs in CSF (miRCURY LNA miRNA Focus PCR Panel, CSF Exosome, Qiagen, YAHS-124Y, 384-well format). qPCR was performed using the miRCURY LNA SYBR Green PCR Kit (Qiagen, 339347) per manufacturer instructions. Any miRNA with a C_q_ value >37 for both antibody and isotype control was excluded from analysis.

### 2.9 Scanning electron microscopy (SEM) and helium ion microscopy (HIM) of EVs on TENPO and eluted EV-MNP complexes

Visualization of EVs immobilized on TENPO was performed via SEM at the Cell & Developmental Biology Microscopy Core at the University of Pennsylvania. Pooled healthy human plasma (Zen-Bio) was preprocessed by thawing at 4°C overnight, keeping the clear phase, and performing a triple spin (at 1,600 x g, 3,000 x g, and 3,000 x g, for 10 minutes each), each time retaining the supernatant. For EV capture, the TENPO protocol was run as described above until after the three washes with 700 μL PBS. Captured EVs were fixed overnight on-chip with 2% glutaraldehyde in 50 mM Na-cacodylate buffer (pH 7.3) at 4°C and washed three times with 50 mM Na-cacodylate buffer using a syringe pump (Braintree Scientific). Membrane sections were cut from the TENPO chip and dehydrated in a graded series of ethanol concentrations through 100% over a period of 1.5 hours.

For SEM/HIM of eluted EV-MNP complexes, EV-MNPs were washed off-chip by flushing devices off-magnet with 300 µL PBS at 50 mL/hr following the conclusion of the three 15 mL/hr 700 µL PBS washes. 30 µL of eluted EV-MNP complexes was then added to glass coverslips (Electron Microscopy Sciences, Cat 72291-02) coated with 100 µg/mL of poly-L-ornithine (Sigma-Aldrich, P3655) and incubated overnight at 4°C. Following removal of the supernatant from the glass coverslips, the coverslips were incubated with 30 µL of 2% glutaraldehyde in 50 mM Na-cacodylate buffer overnight at 4°C. Samples were then washed three times with 200 µL of 50 mM Na-caodylate buffer, with buffer exchange taking place following a 10-minute incubation each time. Dehydration in 100% ethanol was repeated three times.

Following the dehydration steps for both membrane sections and glass coverslips, samples were then incubated for 20 min in 50% hexamethyldisilazane (HMDS) in ethanol followed by three changes of 100% HMDS (Sigma-Aldrich). This was followed by overnight air-drying as described previously.^45^ Samples were stub-mounted and sputter-coated with gold palladium before imaging using a Quanta 250 field emission gun scanning electron microscope (FEI, Hillsboro, OR, USA) at accelerating voltages between 10-30 kV. Additional eluted EV-MNP images were acquired using a Zeiss Orion NanoFab helium-ion microscope at an acceleration of approximately 25,000 kV and a 2 µm field of view.

### 2.10 Cryo-transmission electron microscopy (cryo-TEM) of EVs on TENPO and eluted EV-MNP complexes

For cryo-TEM, to produce sufficient EVs for imaging, TENPO was run as previously described but with an increased starting input volume of 2.5 mL of pooled healthy human plasma (Precision for Medicine). EV-MNPs were washed off-chip by flushing devices off-magnet with 300 µL PBS at 50 mL/hr following the conclusion of the three 15 mL/hr 700 µL PBS washes. EV suspensions were then concentrated using Amicon Ultra-0.5 Centrifugal filters (100K and 30K, Millipore Sigma) before mounting on grids.

300 mesh EM grids (Quantifoil R 1.2/1.3 300 Copper Mesh) were glow discharged for 30 seconds. 3 µL of each EV suspension was deposited onto each grid and vitrified using a Vitrobot Mark IV (Thermo Scientific) set to 100% humidity and 4°C, with a blot time of 4 s. Grids were stored in liquid nitrogen until cassette loading. Imaging was performed using a Glacios Cryo-TEM (Thermo Scientific) operated at 200 kV and visualized at 79,000x magnification.

### 2.11 Isolation of EVs from neuron and astrocyte cell cultures

EVs derived from cell culture were obtained by taking the media from flasks cultured with either mixed cells (neuron and astrocyte) or astrocytes alone. Cells were provided by the Neurons R Us Core at the University of Pennsylvania and were derived from the neocortical tissue of day 18 embryos from Sprague-Dawley rats. Cultures were prepared by mesh-filtering (Crosswire Cloth) cells and resuspending in Minimum Essential Medium (MEM) with Earl’s salts, GlutaMAX (Gibco), 0.6% D-glucose (Sigma-Aldrich), 1% Pen-Strep (Gibco) and 10% Horse Serum (Gibco) for mixed cell culture; or Dulbecco’s Modified Eagle Medium (DMEM, Gibco) with 10% fetal bovine serum (FBS, Sigma-Aldrich) and 1% Pen-Strep for astrocyte cell culture. Cells were then added to T75 flasks (Corning) coated with both 0.08 mg/mL of poly-D-lysine (Sigma-Aldrich) and 0.001 mg/mL of laminin (BD Biosciences) for mixed cell culture, or only 0.08 mg/mL of poly-D-lysine (Sigma-Aldrich) for astrocyte cell culture and allowed to adhere overnight in 37°C at 5% CO_2_. Mixed cell cultures were maintained in B-27 neurobasal medium and 0.4 mM GlutaMAX, and media was collected after 21 days. Astrocyte cultures were maintained in DMEM with FBS and Pen-Strep and passaged at day 21, and media was collected at day 28. Media was frozen at -80°C until TENPO isolation. To prepare cell culture media for TENPO, media was spun at 2,500 x g for 15 minutes. The resulting supernatant was then filtered through a 0.22 µm PES filter (CELLTREAT Scientific Products) before proceeding with the abovementioned TENPO, EV elution, and NTA protocols using 500 µL of input sample volume.

## 3 RESULTS

### 3.1 Study design for isolation and identification of differentially expressed EV miRNAs and proteins

We profiled neuron- and astrocyte-derived EV miRNAs and circulating plasma proteins from 137 patients across five pathologically confirmed disease cohorts: LBD, AD, AD/LBD, AD/ALB, and controls (**Fig. 1A**). Patient numbers per cohort and average ages for symptom onset, sample collection, and death, as well as additional demographic, clinical, and neuropathologic characteristics are summarized in **Table 1**.

**Figure 1:**
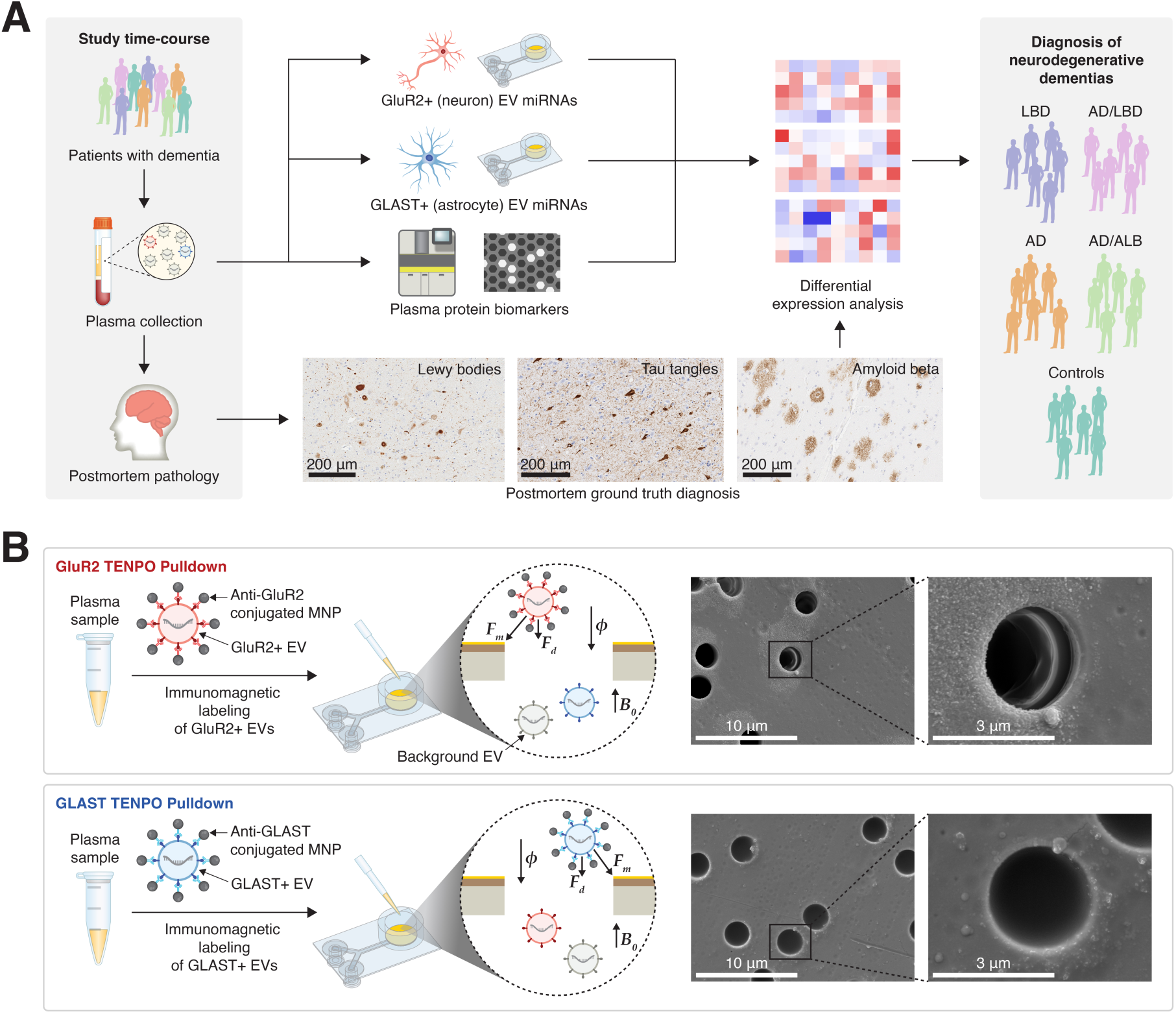
Outline of the study and differential classifications performed. (A) Outline of study time-course and sample processing for EVs. Antemortem plasma samples with postmortem pathology confirmation of neurological diagnoses were processed to isolate tissue-specific EVs via our microfluidic platform, alongside plasma protein biomarkers via commercial digital ELISA, for patients with AD, LBD, AD/LBD, AD/ALB and controls. (B) Left: schematic of operation of TENPO platform for tissue-specific EV isolation using antibody-labeled MNPs; right: SEM images of EVs isolated via GluR2 (top) and GLAST (bottom) pulldowns immobilized on TENPO. Abbreviations: LBD, Lewy body disease; AD, Alzheimer’s disease; AD/LBD, mixed Alzheimer’s and Lewy body disease; AD/ALB, Alzheimer’s disease with amygdala Lewy bodies; EV, extracellular vesicle; TENPO, track-etch magnetic nanopore; ELISA, enzyme-linked immunosorbent assay; MNP, magnetic nanoparticle; SEM, scanning electron microscopy.

**TABLE 1.** Demographic, clinical, and neuropathologic characteristics of study participants.

| Characteristic (N = 137) | LBD | AD | AD/LBD | AD/ALB | Controls |
| --- | --- | --- | --- | --- | --- |
| Number of subjects, n (%) | 30 (21.9%) | 31 (22.6%) | 30 (21.9%) | 19 (13.9%) | 27 (19.7%) |
| **Demographic characteristics** |  |  |  |  |  |
| *Sex* |  |  |  |  |  |
| Male, n (%) | 27 (90.0%) | 13 (41.9%) | 18 (60.0%) | 16 (84.2%) | 12 (44.4%) |
| Female, n (%) | 3 (10.0%) | 18 (58.1%) | 12 (40.0%) | 3 (15.8%) | 15 (55.6%) |
| *Race* |  |  |  |  |  |
| White, n (%) | 30 (100%) | 30 (96.8%) | 29 (96.7%) | 19 (100%) | 26 (96.3%) |
| Asian, n (%) | 0 (0%) | 1 (3.2%) | 0 (0%) | 0 (0%) | 0 (0%) |
| White & Asian, n (%) | 0 (0%) | 0 (0%) | 0 (0%) | 0 (0%) | 1 (3.7%) |
| Black, n (%) | 0 (0%) | 0 (0%) | 1 (3.3%) | 0 (0%) | 0 (0%) |
| *Education* |  |  |  |  |  |
| < High school/GED, n (%) | 0 (0%) | 1 (3.2%) | 1 (3.3%) | 1 (5.3%) | 3 (11.1%) |
| High school/GED, n (%) | 18 (60%) | 16 (51.6%) | 14 (46.7%) | 7 (36.8%) | 9 (33.3%) |
| Bachelor's degree, n (%) | 7 (23.3%) | 4 (12.9%) | 7 (23.3%) | 6 (31.6%) | 4 (14.8%) |
| Master's degree, n (%) | 4 (13.3%) | 6 (19.4%) | 5 (16.7%) | 3 (15.8%) | 7 (25.9%) |
| Doctorate, n (%) | 1 (3.3%) | 4 (12.9%) | 3 (10.0%) | 2 (10.5%) | 4 (14.8%) |
| **Clinical characteristics** |  |  |  |  |  |
| Age at sample collection, years* | 74.05 (6.32) | 73.66 (10.01) | 76.69 (9.98) | 77.66 (8.88) | 79.28 (8.71) |
| Age at symptom onset, years*<sup>†</sup> | 69.50 (9.34) | 68.93 (11.88) | 73.54 (13.09) | 73.52 (10.73) | --/-- |
| Age at death, years* | 80.27 (6.99) | 79.37 (10.16) | 83.22 (10.21) | 83.38 (9.96) | 86.57 (9.36) |
| PMI, hours*<sup>‡</sup> | 25 (16) | 22 (15) | 17 (10) | 16 (10) | 24 (14) |
| **Neuropathologic characteristics** |  |  |  |  |  |
| Braak stage<sup>§,¶</sup> | 2 (2, 3) | 6 (5, 6) | 5 (4, 6) | 6 (6, 6) | 2 (2, 3) |
| CERAD score<sup>§</sup> | 1.5 (0, 2) | 3 (3, 3) | 3 (3, 3) | 3 (3, 3) | 0 (0, 1) |
| Thal phase<sup>§</sup> | 3 (2, 4) | 5 (4, 5) | 4 (3.25, 5) | 5 (4, 5) | 1 (0, 3.5) |

Abbreviations: LBD, Lewy body disease; AD, Alzheimer's disease; AD/LBD, mixed Alzheimer's and Lewy body disease; AD/ALB, Alzheimer's disease with amygdala Lewy bodies; CERAD, Consortium to Establish a Registry for Alzheimer's Disease; GED, General Educational Development; PMI, postmortem interval.
\* Data reported as mean (standard deviation).
<sup>†</sup> 2 subjects (n = 2 LBD) did not have data for age at onset.
<sup>‡</sup> 24 subjects (n = 8 AD, n = 8 LBD, n = 6 AD/LBD, n = 1 AD/ALB, n = 1 control) did not have available PMI data.
<sup>§</sup> Data reported as median (lower quartile, upper quartile).
<sup>¶</sup> 7 subjects (n = 2 LBD, n = 4 controls) did not have available Braak stage data.

We compared demographic factors and clinical characteristics between the cohorts and did not observe statistically significant (*P* value < .05 from Chi-square test) differences between each of the five groups in education level or race (**Supplementary Table 1**). We did note statistically significant differences in the sex ratios of the groups, driven by the higher proportion of males (90%) in the LBD cohort and AD/ALB cohort (84.2%) as compared to the AD, AD/LBD, and control groups (**Supplementary Tables 1-2**); however, LBD has previously been shown in the literature to be more common in men.^46^ Statistical comparison of differences via one-way analysis of variance (ANOVA) in mean age of symptom onset, mean age at time of sample collection, and mean age at death yielded no significant *P* value < .05 differences between the five groups (**Supplementary Table 3**).

For the isolation of neuron- and astrocyte-derived EVs from patient plasma, we applied previous workflows from our group on TENPO EV isolation^27^ using GluR2 and GLAST antibodies. On TENPO, only EVs that were sufficiently magnetized (i.e. had enough MNPs bound to targeted surface proteins) were retained on-chip, as these EVs had sufficient magnetophoretic force acting on them to counteract the strong drag forces from fluid flow (**Fig. 1B**). To characterize these TENPO-captured EVs, we performed SEM and HIM for both eluted EV-MNP complexes off-chip and SEM of whole EV-MNP complexes immobilized on-chip; we also performed cryo-TEM of eluted EV-MNPs off-chip to confirm that our immunomagnetic labeling was binding EVs to MNPs. For SEM, we identified particles with consistent size and morphology as EVs within pores of our TENPO devices (**Fig. 1B, Supplementary Fig. 3**). We also identified eluted EVs on SEM, HIM and cryo-TEM bound to MNPs approximately 50 nm in size (**Supplementary Fig. 4, Supplementary Fig. 5**). Eluted EVs were within the typical EV size range, although EVs appeared larger when imaged by cryo-TEM compared to SEM/HIM, consistent with the literature.^47^

To compare the GluR2 and GLAST pulldowns, we also performed NTA on eluted EVs from TENPO and found no differences in EV concentration or size distribution for either the GluR2 or GLAST pulldowns (**Supplementary Fig. 6**). We confirmed the specificity of the GluR2+ EV and GLAST+ EV-derived signals by demonstrating statistically significant increases in expression of multiple brain-relevant miRNAs, when comparing EVs isolated from plasma using GluR2 and GLAST versus isotype controls (**Supplementary Fig. 7**). A commercially available panel for CSF markers (Qiagen) was used for this assay, as it provided a readily accessible measure for brain-specific RNA cargo.

To validate that the GluR2 pulldown isolated neuron-specific EV signals and the GLAST pulldown isolated astrocyte-specific EV signals, we performed TENPO isolation on 500 µL of media obtained from mixed cell cultures (neuron and astrocyte co-culture) or astrocyte-only cell cultures derived from rat neocortical tissue. NTA was performed on EVs eluted off chip, showing additional EV-sized particles isolated using GluR2 TENPO from mixed media that were not present when performing GLAST TENPO on mixed media (**Supplementary Fig. 8A**). Similarly, for astrocyte-only media, NTA showed additional EV-sized particles for GLAST TENPO that were not present for GluR2 TENPO (**Supplementary Fig. 8B**). Furthermore, NTA for GLAST TENPO on mixed media and GluR2 on astrocyte-only media showed similar size distributions to bead-only NTA measurements, indicating that the particles detected on NTA for these two conditions may be primarily from beads that have passed into the eluate (**Supplementary Fig. 8C**), with a slight rightward shift of the NTA peak for the former due to the presence of astrocytes within the mixed cultures.

### 3.2 Neuron- and astrocyte-derived EV miRNAs and plasma proteins differ in expression between LBD and AD

The TENPO-isolated neuron and astrocyte EV miRNAs alongside the plasma protein measurements defined a biomarker expression profile for each patient in this study. In a comparison of subjects with LBD (n = 30) versus those with AD (n = 31), in which each subject did not exhibit any degree of mixed postmortem pathology of the two diseases, we observed multiple nucleic acid and protein biomarkers which showed significant differential expression after FDR correction. To visualize the spread of miRNA and protein expression across our LBD and AD cohorts, we performed hierarchical clustering of the z-score of log_2_(expression) data between patients (**Fig. 2A**). We identified 16 differentially expressed miRNAs from GluR2+ EVs (11 upregulated, five downregulated), eight differentially expressed miRNAs from GLAST+ EVs (seven upregulated, one downregulated), and four differentially expressed plasma protein biomarkers, including Aβ42/Aβ40 ratio (one upregulated, three downregulated) (**Fig. 2B**).

**Figure 2:**
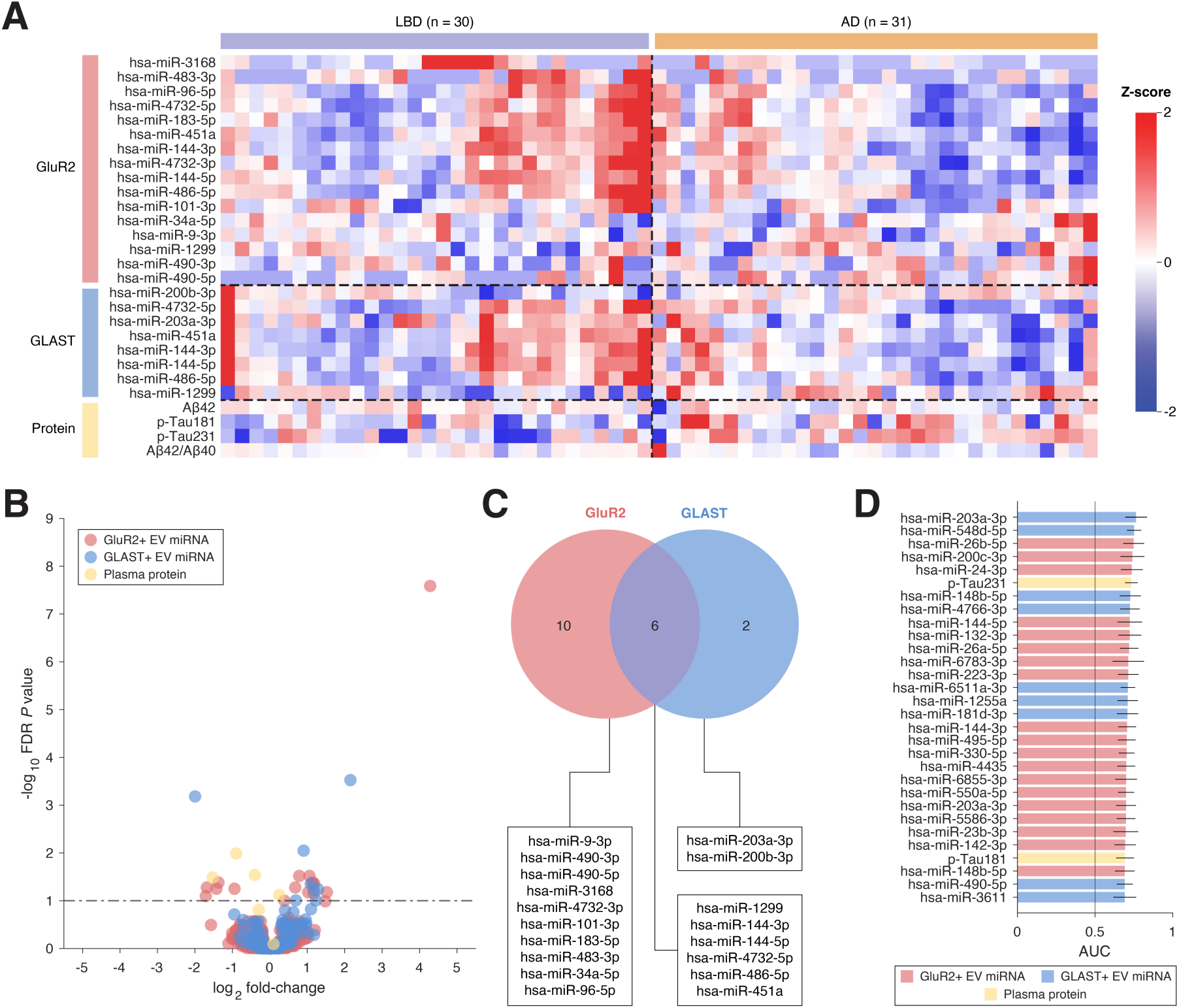
Differential expression of GluR2+ EV miRNAs, GLAST+ EV miRNAs, and plasma proteins in the LBD versus AD comparison. (A) Heatmap of z-score of log_2_(expression) for biomarkers with FDR-corrected *P* value < .1. Subjects (columns) are hierarchically clustered within cohort and biomarkers within each compartment (rows) are sorted by descending fold-change. (B) Volcano plot demonstrating differential expression of GluR2+ EV miRNAs, GLAST+ EV miRNAs, and plasma proteins. (C) Venn diagram showing overlap in FDR *P* value significant miRNAs between GluR2+ EVs and GLAST+ EVs. (D) Top 30 biomarkers in all compartments ranked by descending AUC. Error bars represent standard error from bootstrapping 10x. Abbreviations: LBD, Lewy body disease; AD, Alzheimer’s disease; EV, extracellular vesicle; FDR, false discovery rate; AUC, area under the curve.

To assess the extent to which the GluR2 versus GLAST compartments yielded complementary biological information, we identified the overlap between differentially expressed miRNA markers from each pulldown. There were six overlapping markers, each of which exhibited upregulation or downregulation within both compartments: miR-144-3p, miR-144-5p, miR-4732-5p, miR-486-5p, and miR-451a were upregulated in both compartments; and miR-1299 was downregulated in both compartments (**Fig. 2C**). We also identified a combined 12 differentially expressed markers (10 in GluR2, two in GLAST) unique to a single pulldown, highlighting both shared and distinct molecular signatures among the different EV subtypes.

We assessed the capability of each individual nucleic acid and protein marker to classify LBD versus AD by performing a ROC analysis and reporting the AUC; the AUCs for the top 30 biomarkers are shown in **Fig. 2D**. A total of five markers in the top 30 markers ranked by AUC was both statistically significant for differential expression after FDR correction and had a significant AUC compared to a non-predictive (AUC = 0.5) marker via Delong’s test. Of these markers, GLAST+ EV miR-203a-3p had the greatest AUC at 0.76 (Delong’s *P* value = .0094), followed by p-Tau231 (AUC = 0.73, Delong’s *P* value = .0048), GluR2+ EV miR-144-5p (AUC = 0.72, Delong’s *P* value = .026), GluR2+ EV miR-144-3p (AUC = 0.71, Delong’s *P* value = .045), and p-Tau181 (AUC = 0.69, Delong’s *P* value = .039).

The miRNA and protein profiles revealed complex patterns of expression, with outliers and heterogeneity within disease cohorts, even for strongly predictive markers (e.g. GluR2+ EV miR-144-5p) (**Supplementary Fig. 9**). Notably, for the markers upregulated in LBD versus AD, expression was highest in subclusters of patients in both the GluR2 and GLAST pulldowns (**Fig. 2A**). Some downregulated GluR2 markers (miR-490-3p, miR-34a-5p, miR-1299, miR-9-3p) appeared to be influenced by outlier expression profiles in a few patients, resulting in correspondingly low AUCs (AUCs of 0.58, 0.55, 0.54 and 0.53, respectively) (**Fig. 2D**, **9A**). A single outlier patient within the AD cohort had high expression in the Aβ42/Aβ40 ratio, driving overall downregulation of Aβ42/Aβ40 and yielding an AUC (AUC = 0.68, Delong’s *P* value = .12) that was not significant. The ROC curves for the Delong’s-significant differentially expressed markers for LBD versus AD are shown in **Supplementary Fig. 10A-C**. We also performed a comparison of the AUCs for the top 30 miRNA markers between the GluR2 and GLAST pulldowns to assess their degree of correlation, identifying that the pulldowns did not yield significant correlation for these miRNAs (**Supplementary Fig. 10D**).

### 3.3 Dementia patients exhibit altered miRNA and protein profiles relative to controls

We compared the biomarker profiles of patients with pathologically defined dementia to clinically normal controls using three comparisons: (1) a comparison of combined dementias (n = 30 LBD, n = 31 AD) versus n = 27 controls, (2) a comparison of n = 31 AD patients versus n = 27 controls, and (3) a comparison of n = 30 LBD patients versus n = 27 controls. As performed previously for LBD versus AD, we generated hierarchically clustered heatmaps of the z-score of log_2_(expression) between patients for all three comparisons (**Supplementary Fig. 11A, D, and G**).

For LBD and AD versus control, we identified a total of seven differentially expressed miRNAs (two GluR2, five GLAST, with one overlapping miRNA between the pulldowns) and two differentially expressed protein biomarkers (Aβ42 and Aβ42/Aβ40). Of these biomarkers, three GLAST+ EV miRNAs and one protein were upregulated in the combined dementia cohort, while the remaining biomarkers were upregulated in the control cohort (**Supplementary Fig. 11B, 12A-C**). By performing an ROC analysis on each biomarker with a statistically significant fold-change (**Supplementary Fig. 13A-B**), we identified one GLAST+ EV miRNA (miR-204-5p, AUC = 0.67, Delong’s *P* value = .062) and two protein biomarkers (Aβ42/Aβ40, AUC = 0.67, Delong’s *P* value = .051; Aβ42, AUC = 0.66, Delong’s *P* value = .045) with an AUC significant compared to a non-predictive (AUC = 0.5) marker via Delong’s test, for which Aβ42/Aβ40 featured the highest classifying AUC (**Supplementary Fig. 11C**). miR-204-5p has been previously identified as a dementia biomarker in CSF; however, it was downregulated in AD patients versus controls,^48^ whereas it is upregulated in the combined LBD and AD cohort in this study.

In our comparison of AD versus control, we identified a total of 13 miRNAs (11 GluR2, two GLAST, with one overlapping miRNA between the pulldowns) and three proteins (p-Tau181, Aβ42, and Aβ42/Aβ40) which showed statistically significant differences between AD versus control. Within this group of biomarkers, eight miRNAs (seven from GluR2, one from GLAST), Aβ42/Aβ40, and p-Tau181 were upregulated in the AD cohort; the remaining biomarkers were upregulated in the control cohort (**Supplementary Fig. 11E, 12D-F**). ROC analysis of these biomarkers yielded four biomarkers (GluR2+ EV miR-132-3p alongside proteins p-Tau181, Aβ42, and Aβ42/Aβ40) with statistically significant AUCs per Delong’s test (Delong’s *P* value = .0039, .027, .017, and .018, respectively), with miR-132-3p having the highest AUC of the group at 0.76 (**Supplementary Fig. 11F, 13C-D**). Aβ42 was upregulated in controls in a manner consistent with the literature.^19, 49^ p-Tau181 was also upregulated in AD versus controls, similar to what has been previously reported in the literature.^4^ While overall Aβ42/Aβ40 expression was upregulated for both the combined dementia versus control and AD versus control comparisons, which runs contrary to the literature,^19, 49^ this was largely driven by one outlier, as median Aβ42/Aβ40 expression for the combined dementia and AD cohort were lower than for the control group.

Our comparison of LBD versus control yielded only two differentially expressed biomarkers (p-Tau231 and GLAST+ EV miR-200b-3p) (**Supplementary Fig. 11H, 12G-H**), and only p-Tau231 had a statistically significant AUC at 0.69 (Delong’s *P* value = .020) (**Supplementary Fig. 11I, 13E)**. The limited number of differentially expressed markers between LBD versus control could explain the decrease in identified markers for combined dementia versus control as compared to AD versus control and is consistent with the documented lack of diagnostic biomarkers for LBD to date.

### 3.4 EV miRNA biological function and pathway analysis yields insights into mechanistic underpinnings

To determine whether the identified biomarkers relate to known pathways in the biology of LBD and AD, we performed Gene Ontology (GO) and Kyoto Encyclopedia of Genes and Genomes (KEGG) analyses on the union of differentially expressed miRNAs from each individual EV pulldown. We pinpointed the top 10 terms per GO category [biological process (BP), cellular component (CC), and molecular function (MF)] (**Fig. 3A, 3C**) and top 10 KEGG pathways (**Fig. 3B, 3D**) by the number of target genes affected by each set of miRNAs. Additional significantly enriched (FDR *P* value < .05) GO BP terms associated with brain-related processes and neuroinflammation are shown in **Supplementary Table 4** and significantly enriched (FDR *P* value < .05) KEGG pathways associated with the nervous system, neuron development, and brain-specific diseases are shown in **Supplementary Table 5**. Within the GluR2 pulldown, a maximum of 14 miRNAs were identified as being involved in any given GO term or KEGG pathway. Likewise, within the GLAST pulldown, our GO and KEGG analyses identified terms involving a maximum of seven miRNAs.

**Figure 3:**
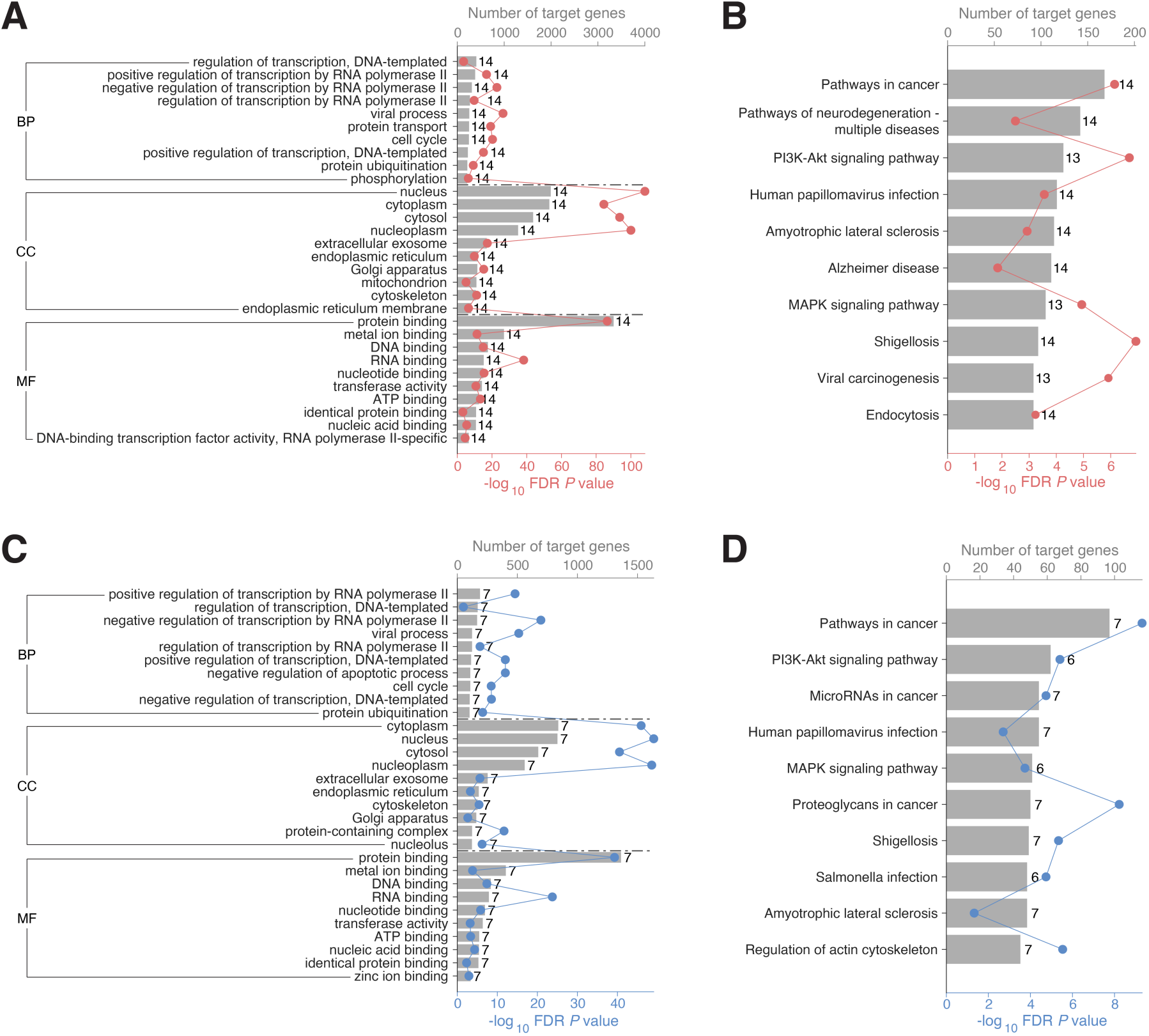
GO and KEGG analysis of FDR *P* value significant miRNAs in the LBD versus AD comparison. GO and KEGG pathway analyses were performed via DIANA miRPath v4.0 using the TarBase v8.0 database. FDR *P* values were calculated using a one-sided Fisher’s exact test. The top 10 (ranked by number of target genes) terms within each of the three GO categories (BP, CC, MF) and top 10 (ranked by number of target genes) KEGG pathways were identified for each pulldown. (A) Top 10 terms within each GO category for GluR2+ EV miRNAs. (B) Top 10 KEGG pathways for GluR2+ EV miRNAs. (C) Top 10 terms within each GO category for GLAST+ EV miRNAs. (D) Top 10 KEGG pathways for GLAST+ EV miRNAs. In all panels, each bar is labeled to the right with the number of differentially expressed miRNAs associated with the given GO term or KEGG pathway. Abbreviations: LBD, Lewy body disease; AD, Alzheimer’s disease; EV, extracellular vesicle; FDR, false discovery rate; GO, Gene Ontology; KEGG, Kyoto Encyclopedia of Genes and Genomes; BP, biological process; CC, cellular component; MF, molecular function.

The GO and KEGG terms identified in the present study were both directly and indirectly linked to neurodegeneration and neuroinflammation, but also included terms including “cell cycle” and “regulation of transcription” (GO BP) that are shown to be dysregulated in studies concerning general aging or other age-related pathologies such as cancer.^50^ Of note, “negative regulation of apoptotic process” only appeared as a BP term within the GLAST pulldown. While LBD and AD are marked by neuronal cell death,^51^ they are also associated with upregulation in both astrocytic pro-inflammatory or neuroprotective, proliferative states,^52^ although response appears to be specific to both brain region and disease stage.^53^

The KEGG analysis revealed significant enrichment in multiple neurodegenerative pathways and diseases. The MAPK signaling and PI3K-Akt pathways, which appeared for both pulldowns, are known regulators of synaptic plasticity and neuronal survival.^54, 55^ Additionally, “endocytosis”, which appeared as a GluR2 KEGG-enriched pathway, is consistent with our understanding of uptake and spread of pathological proteins in neurodegeneration.^56^ Lastly, “regulation of the actin cytoskeleton”, enriched within the GLAST pulldown, maintains normal synaptic function and is impaired in both LBD and AD.^57^

miRNAs control gene expression by binding to target mRNAs; therefore, the examination of gene targets influenced by differential miRNA expression can demonstrate the functional implications of miRNA dysregulation in driving disease states. The top 10 enriched KEGG pathways showed the greatest interaction between differentially expressed GluR2+ EV miRNAs with target genes *IGF1R* (inhibition of which is linked to Aβ accumulation and tau phosphorylation),^58^ *KIF5B* (inhibition of which causes memory deficits in mice),^59^ and *MAPK1* (which controls a wide network of genes affected in LBD and AD)^60, 61^ (**Supplementary Fig. 14A**). For differentially expressed GLAST+ EV miRNAs, KEGG pathways showed the greatest interaction with apoptosis regulators *BCL2*^62^ and *MYC*,^63^ along with *PTK2*, which controls tau-induced neurotoxicity^64^ (**Supplementary Fig. 14B**).

### 3.5 Feature selection identifies multimodal biomarker panels for differentiating LBD from AD

We used LASSO feature selection to identify candidate miRNAs and proteins which were most informative for classifying LBD versus AD when in combined panels. For input into LASSO, we used all neuron- and astrocyte-derived EV miRNAs which cleared the expression threshold for the Qiagen RNA sequencing analysis pathway to generate an FDR-corrected *P* value. Similarly, we included all plasma proteins that generated an FDR-corrected *P* value.

In sweeping the number of selected biomarkers from two to 20, we generated a composite list of biomarkers including 14 GluR2+ EV miRNAs, four GLAST+ EV miRNAs, and the plasma proteins p-Tau231 and p-Tau181 (**Table 2**), with many of the LASSO-selected markers having been identified by previous literature concerning neurological conditions (**Supplementary Table 6**). As with our previous analyses, we generated a heatmap of the z-score of log_2_(expression) of LASSO-selected biomarkers with hierarchically clustered patients (**Fig. 4A**). We also performed a pairwise Kendall tau correlation analysis between each of the biomarkers (**Fig. 4B**), identifying only limited correlation. While several biomarkers did not feature statistically significant differential expression between LBD versus AD, the top 14 biomarkers ranked by AUC, in addition to GluR2+ EV hsa-miR-4748, were all found to have AUCs significantly above 0.5 per Delong’s test.

**Figure 4:**
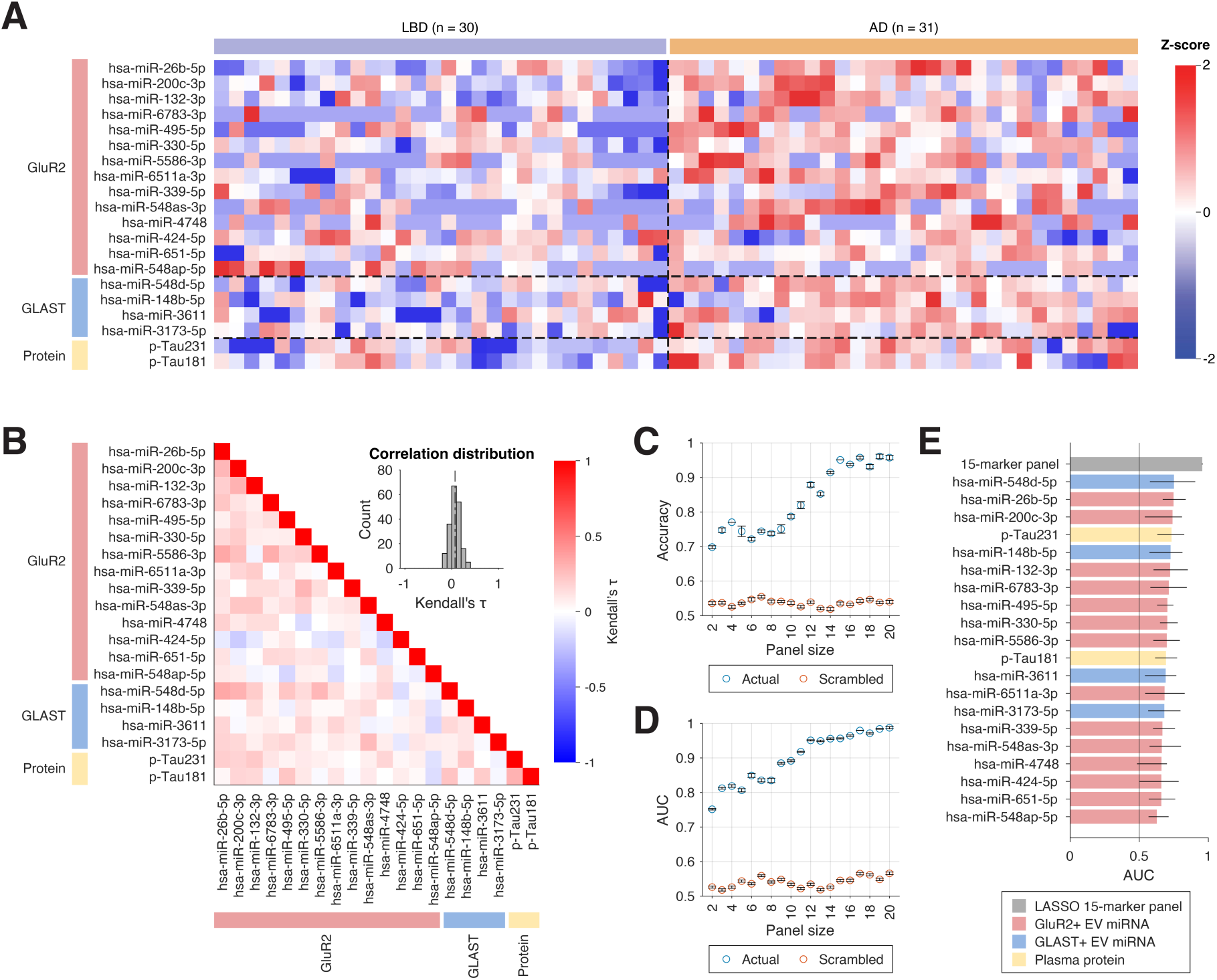
LASSO analysis for differentiating LBD (n = 30) from AD (n = 31). **(A)** (A) Heatmap of z-score of log_2_(expression) for LASSO-selected biomarkers. Subjects (columns) are hierarchically clustered within cohort and biomarkers within each compartment (rows) are sorted by descending AUC. (B) Kendall correlation staircase plots identifying the extent to which biomarker information was correlated between the LASSO-selected GluR2+ EV, GLAST+ EV, and protein biomarkers. Biomarkers are sorted within compartments by AUC. The inset shows the correlation distribution of Kendall’s τ, where the dotted line represents the median count. (C) LASSO panel accuracy versus panel size for classifying LBD versus AD, shown in blue; accuracy is assessed via 10-fold cross-validation, with error bars from 5 repeats of panel training on the LBD vs AD patient groups. Average accuracy and standard error for control experiments performed by scrambling patient labels 10x are shown in orange. (D) LASSO panel AUC versus panel size for classifying LBD versus AD, shown in blue with error bars as described in (C). Average AUC and standard error for the same control experiments described in (C) are shown in orange. (E) AUCs for the 15-marker LASSO panel and individual LASSO biomarkers, sorted by descending AUC. Error bars represent 95% confidence intervals, calculated from 5x repeats of 10-fold cross validation for the 15-marker panel or from bootstrapping 10x for individual markers. Abbreviations: LBD, Lewy body disease; AD, Alzheimer’s disease; EV, extracellular vesicle; LASSO, least absolute shrinkage and selection operator; AUC, area under the curve.

**TABLE 2.** Nucleic acid and protein biomarkers selected via LASSO for comparing LBD versus AD for panel sizes 2-20. The table shows the incremental addition of biomarkers as the panel size increases. For each panel size, *n*, the marker newly added to the panel is listed. Panels of size *n* include this newly added marker in addition to all markers from panel sizes < *n.* Markers are annotated with their associated compartment (GluR2, GLAST, or protein) in parentheses.

| Panel size ($n$) | Newly added marker (at size $n$) |
| --- | --- |
| 2 | hsa-miR-548d-5p (GLAST), hsa-miR-26b-5p (GluR2) |
| 3 | p-Tau231 (protein) |
| 4 | hsa-miR-200c-3p (GluR2) |
| 5 | hsa-miR-330-5p (GluR2) |
| 6 | hsa-miR-6511a-3p (GluR2) |
| 7 | hsa-miR-495-5p (GluR2) |
| 8 | hsa-miR-132-3p (GluR2) |
| 9 | hsa-miR-148b-5p (GLAST) |
| 10 | hsa-miR-339-5p (GluR2) |
| 11 | hsa-miR-4748 (GluR2) |
| 12 | hsa-miR-3611 (GLAST) |
| 13 | hsa-miR-548as-3p (GluR2) |
| 14 | hsa-miR-3173-5p (GLAST) |
| 15 | hsa-miR-5586-3p (GluR2) |
| 16 | p-Tau181 (protein) |
| 17 | hsa-miR-424-5p (GluR2) |
| 18 | hsa-miR-6783-3p (GluR2) |
| 19 | hsa-miR-548ap-5p (GluR2) |
| 20 | hsa-miR-651-5p (GluR2) |

Abbreviations: LASSO, least absolute shrinkage and selection operator; LBD, Lewy body disease; AD, Alzheimer's disease.

To assess the combined performance of biomarkers within each panel, we used an ensemble model on the entire cohort of LBD versus AD patients with 10-fold cross-validation repeated 5x, displaying a plateau in panel accuracy for a panel size of 15 (accuracy = 0.95) (**Fig. 4C**), at which the panel AUC = 0.96 (**Fig. 4D**). The ROC curve for the combined 15-marker panel shows that the panel greatly outperformed each individual LASSO-selected biomarker (**Supplementary Fig. 15**), showing an improvement in AUC of 0.21 over the best performing individual LASSO biomarker, and non-overlapping 95% confidence intervals between the 15-marker panel and any of the individual LASSO biomarkers (**Fig. 4E**).

We compared the model’s performance to control experiments in which LASSO was implemented for patients whose labels were randomly scrambled using MATLAB’s random permutation function. The resulting panels were assessed for diagnostic utility in categorizing patients based on their true labels. This was repeated 10 times with the average accuracy and AUC falling below 0.6 for all panel sizes (**Fig. 4C, 4D**). At all panel sizes, the cross-validated panel AUCs were also significantly higher than for their associated scrambled controls *(P* value < .001 via the Mann-Whitney U test).

We then examined the relative expression levels of LASSO-selected biomarkers in mixed-pathology patients with histopathologic diagnoses of AD/LBD (n = 30) or AD/ALB (n = 19) as compared to LBD and AD patients (**Supplementary Fig. 16**). We observed that the AD/LBD and AD/ALB patient cohorts generally featured median biomarker expression levels in between the LBD and AD groups, which is consistent with the status of AD/LBD and AD/ALB as mixed pathologies. For most miRNA biomarkers we examined, we did not observe large differences between the AD/LBD and AD/ALB cohorts; however, in both LASSO-selected protein biomarkers p-Tau181 and p-Tau231, we observed higher expression of these proteins in AD/ALB versus AD/LBD patients.

## 4 DISCUSSION

While clarifying the role of EV cargoes within LBD and AD needs further work, this study identifies biomarkers that may offer new avenues for diagnostic and therapeutic strategies in these two diseases, and supports the hypothesis that EVs are linked to signaling changes associated with neurodegeneration. For example, β-amyloid plaques within the AD brain are interdigitated by EVs,^65^ suggesting that they participate in plaque formation, and EVs are posited to propagate neurotoxic Aβ, tau, and α-syn between astrocytes and neurons,^25^ leading to increased neuronal death and proximal microglial activation.^66^ Astrocyte-derived EVs that cross the BBB can also increase neuroinflammation through peripheral leukocytic recruitment into the brain.^66^ However, EVs may still retain some role in neuroprotection within the context of dementia; in the AD brain, EV-bound PrP^c^ preferentially binds Aβ and promotes its fibrillization and decreased toxicity.^25^ As the molecular content of EVs is influenced by their originating cell’s pathological state,^24^ our ability to capture signal from these two specific cell types allows us to restrict our view to biomarkers that are specific to the disease and may be involved these pathways.

The heterogeneity in individual miRNAs and proteins across our patient cohorts underscores the difficulty of relying on a single marker for differential diagnosis, as shown by the modest AUCs for individual miRNAs. In contrast, the integration of a minimally correlated multi-marker panel selected by LASSO from all three compartments reveals an identifiable pattern of heatmap expression and results in clinically relevant (AUC > 0.8) AUCs for all panel sizes of at least three biomarkers, and high AUCs (AUC > 0.9) for panel sizes > 10. To date, even gold-standard AD biomarkers such as p-tau217 have only achieved an AUC of 0.84 in distinguishing AD for patients with early-onset/atypical dementia.^16^ Similarly, α-syn seed amplification assays (SAAs), which have emerged as a method for LBD diagnosis, have primarily achieved success in CSF, brain tissue homogenates,^67^ and skin.^68^ While preliminary SAA results from the serum of non-autopsy-confirmed subjects showed good performance for PD versus controls (AUC = 0.96 initial cohort; AUC = 0.86 in blinded independent cohort), notably, 4 of 25 AD subjects were SAA-positive.^69^

Our AUCs suggest strong predictive performance of our LASSO-identified biomarker panel. This study utilizes exceptionally valuable biospecimens, which can only be obtained postmortem; to mitigate this constraint, and given the confounding effect of combining non-pathology-confirmed samples with this cohort, the AUCs and accuracies in our approach were generated using a robust cross-validation strategy. This allowed us to perform an internal assessment of model performance, as well as maximize our sample size for biomarker discovery. Although panels larger than 10 markers may seem considerable, the ability to measure large biomarker panels has become more accessible with the advancement of multiplexed technologies, which may be employed in future validation experiments with an independent cohort. This larger panel size may also facilitate reproducibility across larger cohorts, mitigating the variability inherent in dementia populations.

To date, histopathology remains the gold standard for the diagnosis of LBD, AD, and mixed-pathologies.^3, 70^ The inability to reliably detect non-AD or mixed pathologies prior to death remains an obstacle in designing effective clinical trials, particularly given that postmortem studies indicate 35-50% of PD patients and >70% of DLB patients exhibit AD co-pathology.^70^ Clinically, AD/LBD and AD/ALB bear close symptomatic resemblance to pure AD patients in advanced stages.^51^ Existing single LBD and AD biomarkers, like SAAs and p-tau217, may not be effective in detecting these mixed pathology cases; recent studies indicate that SAAs may still produce a positive result for some patients with Lewy bodies that do not meet neuropathological criteria for LBD^17, 18^ and p-Tau217 is unable to predict presence or absence of LBD co-pathology in AD patients.^4^ Likewise, imaging biomarkers such as ^18^F-fluorodeoxyglucose-positron emission tomography (^18^F-FDG PET) have shown limited performance in distinguishing between AD with concomitant amygdala-predominant versus limbic/neocortical Lewy bodies.^71^ While we observed some overlap in individual LASSO-selected biomarker profiles between pure and mixed pathology cohorts, select markers showed identifiable trends in median expression level associated with degree of Lewy pathology.

Although the sample size limited our statistical power, these findings, and the lack of reliable single biomarkers for detection of mixed pathology thus far, underscore the benefits of using larger multimodal panels to increase the likelihood of stratifying varying Lewy body and AD co-pathology. This remains a critical challenge, as not knowing the true state of enrolled participants would affect the design of clinical trials in two ways: (1) in clinical trials aimed at targeting a single disease, the estimated rates of decline and survival times would be inappropriate for patients with hidden co-pathology, impacting measures of trial success^72^ and (2) because the anticipated mechanisms of decline would be improperly characterized, there is the potential for drug inefficacy or toxicity induced by off-target activity.^73^ Conversely, an attempt to exclude mixed pathology patients could result in the development of therapeutics that may not work on the general population. Future studies involving patients along the mixed AD and LBD pathological spectrum will need to identify means of understanding the biology driving the disease process to better characterize the “ground truth” for determining differentially expressed biomarkers between these populations.

There are several limitations of our study and opportunities for future work. First, it is important to note that our biomarker measurements are collected at a time point separated by years from the postmortem pathology. While detectable cognitive changes were present in our patient populations, we cannot determine the precise point at which each patient was located within the course of their disease. This uncertainty could have affected subjects with mixed pathology (AD/LBD and AD/ALB), as patients may not have had both diseases at sample draw or may have lacked sufficient core clinical features to detect one or both diseases. Furthermore, each patient had a recorded clinical diagnosis at the time of visit; however, given the number of subjects with a diagnosis of mild cognitive impairment (MCI), the potential for non-AD diagnoses including MCI to evolve into AD, and the lack of knowledge of when patient MCI evolved into frank dementia, we did not attempt to directly correlate accuracy of clinical diagnosis versus histopathologic diagnosis.

Second, given the sample type (i.e. plasma collected years before death for patients with pathology-proven subtypes of dementia), our sample size, subject diversity, and sample volumes were limited. Because our plasma samples were collected as part of the ADRC or MCSA studies, our subjects reflect the predominantly white population in the upper Midwest. Likewise, we had a high percentage of males within both LBD and AD/ALB cohorts; while this is consistent with the literature for LBD,^46, 74^ for AD/ALB, this discrepancy may have due to the limited cohort size. Recent studies have shown that both race^75^ and sex differences^46, 74^ may play important roles in disease progression, clinical presentation, and survival of LBD, AD, and mixed pathology patients; thus, further studies with greater diversity and balanced sex from multiple collection centers are needed to validate these markers within the general population. Finally, since our study focuses on detecting biomarkers within a complex aging population, many of the subjects had concomitant non-LBD and non-AD pathologies, including vascular dementia, primary age-related tauopathy, and age-related degeneration, that may have also driven miRNA and protein expression.

Future work aims to address these limitations by using the set of biomarkers identified in this study in multi-omic and multi-center clinical studies to survey neurodegenerative dementia across more diverse patient populations. This study also identifies a set of candidate biomarkers which could be interrogated in cellular models of neurons and astrocytes to further the mechanistic understanding of the biology of LBD and AD. Given the accelerating pace of the development of therapeutics in neurodegenerative dementia, this work articulates a multimodal and biomarker-driven approach to providing greater diagnostic clarity for providers and patients in understanding and treating these complex diseases.

## ACKNOWLEDGMENTS

The authors thank Drs. Yangzhu Du and Honghong Sun of the Human Immunology Core at the Perelman School of Medicine at the University of Pennsylvania for assistance with digital ELISA assays. The authors also would like to thank Dr. Yuri Veklich, Dr. Jamie Ford, and Neha Srikumar for assistance with imaging of EVs via scanning electron microscopy and helium ion microscopy; Dr. Vera Moiseenkova-Bell, Dr. Dr. Prerana Gogoi, and Dr. Ruth Anne Pumroy for assistance with imaging EVs via transmission electron microscopy; Dr. Luca Musante for assistance with performing nanoparticle tracking analysis; Kyra Kovacic and Dr. Jonathan Schug for assistance with RNA sequencing; Dr. Josh Buser for assistance in fabrication and assembly of TENPO chips; Dr. Kryshawna Beard for assistance in conducting cell culture; Dr. Hiroaki Sekiya for taking representative pathology pictures for this manuscript; and Dr. Fabrice Lucien-Matteoni for sample storage. The authors particularly thank all participants, volunteers, and coordinators who contributed to this research.

## DATA AVAILABILITY

The data that support the findings of this study are available from the corresponding author, upon reasonable request.

## CONSENT STATEMENT

All participants or their proxies provided informed consent for antemortem sample collection and postmortem analysis. All methods were conducted in accordance with appropriate clinical guidelines and recommendations, and were approved by the Mayo Clinic IRB (IRB# 712-98 for ADRC; 14-004401 for MCSA). Confidentiality of patient information was maintained throughout the study.

