## Supplementary Material for "Brain-derived extracellular vesicle microRNAs in Lewy body and Alzheimer’s disease"

**Supplementary Fig. 1. Analysis of batch-to-batch variation between sequencing runs.** Left: 10 libraries were rerun between two different sequencing runs and the Kendall correlation of their reads was assessed. As a control, the correlation of each library with 10 randomly chosen libraries from the same sequencing round was also assessed (error bars from n = 5 iterations). Right: example scatter-plot demonstrating close correlation of CPM-normalized reads between two different sequencing runs of the same patient sample. Abbreviations: CPM, counts-per-million.

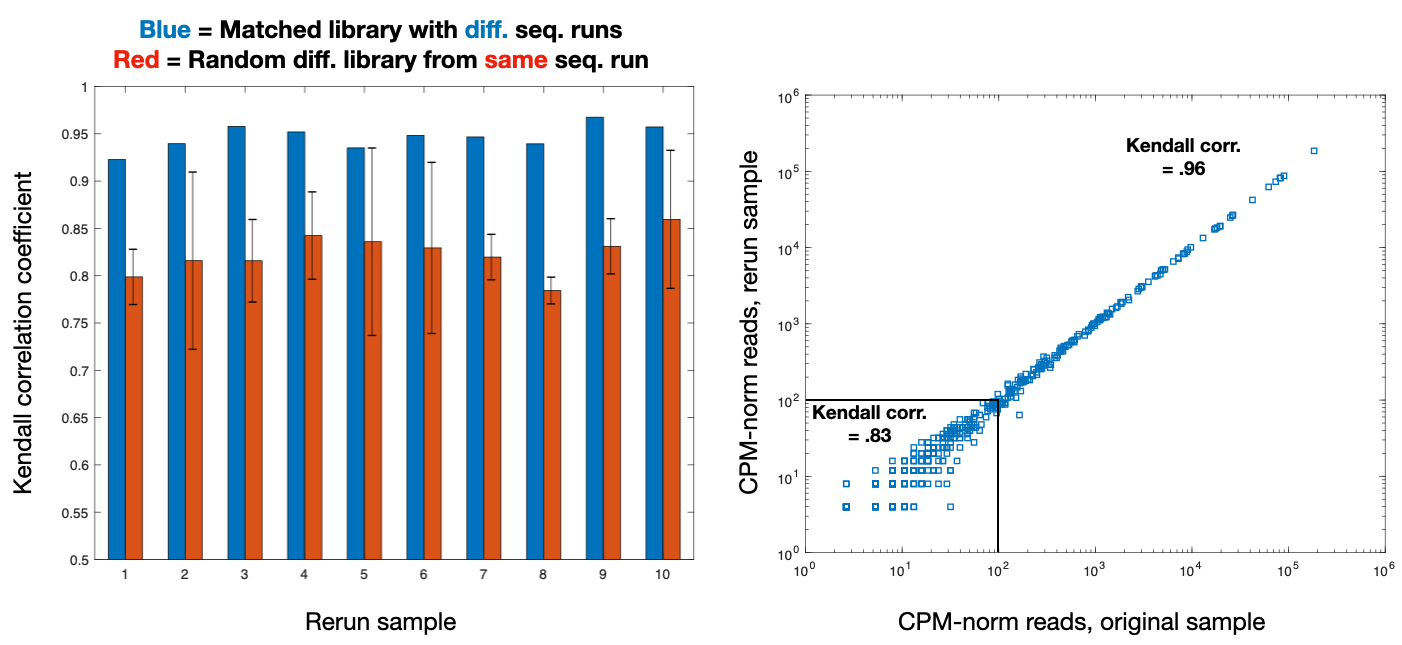

**Supplementary Fig. 2. Analysis of impact of batch-to-batch variation between sequencing runs on relative expression of miRNAs between patients.** Example data shown for higher-read (left) and lower-read (right) nucleic acid marker. Abbreviations: CPM, counts-per-million.

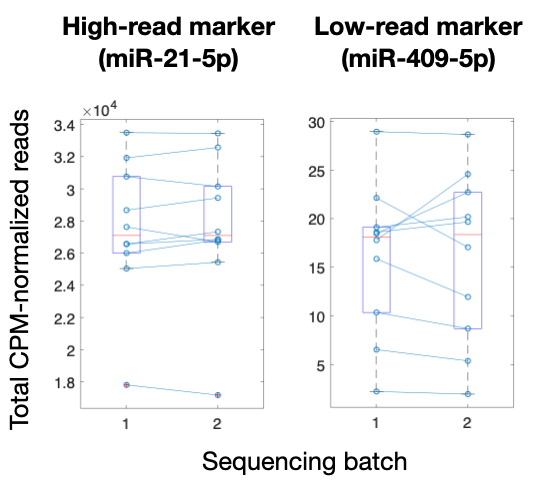

**Supplementary Table 1. Table of grouped Chi-square tests for sex, education, race.** Chi-square tests were performed across the five groups compared in this study (AD, LBD, AD/LBD, AD/ALB, and controls). Abbreviations: LBD, Lewy body disease; AD, Alzheimer’s disease; AD/LBD, mixed Alzheimer’s and Lewy body disease; AD/ALB, Alzheimer’s disease with amygdala Lewy bodies.

| **Comparison** | ***P* value** |
| --- | --- |
| Sex | 1.270*10^-4^ |
| Education | 0.652 |
| Race | 0.520 |

**Supplementary Table 2. Table of individual Chi-squares for sex.** Chi-square tests were performed to determine significance in differences observed in sex ratios between the patient groups. Abbreviations: LBD, Lewy body disease; AD, Alzheimer’s disease; AD/LBD, mixed Alzheimer’s and Lewy body disease; AD/ALB, Alzheimer’s disease with amygdala Lewy bodies.

| **Comparison** | ***P* value** |
| --- | --- |
| AD vs. LBD | 7.817*10^-5^ |
| AD vs. AD/LBD | 0.158 |
| AD vs. AD/ALB | 4.899*10^-6^ |
| AD vs. control | 0.427 |
| LBD vs. AD/LBD | 0.0061 |
| LBD vs. AD/ALB | 0.0015 |
| LBD vs. control | 4.419*10^-5^ |
| AD/LBD vs. AD/ALB | 0.0001 |
| AD/LBD vs. control | 0.105 |
| AD/ALB vs. control | 1.170*10^-7^ |

**Supplementary Table 3. Table of one-way ANOVAs for numeric demographic/clinical factors.** One-way ANOVAs were performed across the five groups compared in this study (AD, LBD, AD/LBD, AD/ALB, and controls) unless otherwise indicated. Abbreviations: ANOVA, analysis of variance; LBD, Lewy body disease; AD, Alzheimer’s disease; AD/LBD, mixed Alzheimer’s and Lewy body disease; AD/ALB, Alzheimer’s disease with amygdala Lewy bodies; PMI, postmortem interval.

| **Comparison** | ***P* value** |
| --- | --- |
| Age at death | 0.0696 |
| Age at collection | 0.1628 |
| Age at symptom onset (AD vs. LBD vs. AD/LBD vs. AD/ALB) | 0.2827 |
| PMI | 0.1343 |

**Supplementary Fig. 3. SEM micrographs of EVs isolated on chip from healthy human plasma for GluR2 (A) and GLAST (B) EV pulldowns.** Abbreviations: SEM, scanning electron microscopy; EV, extracellular vesicle.

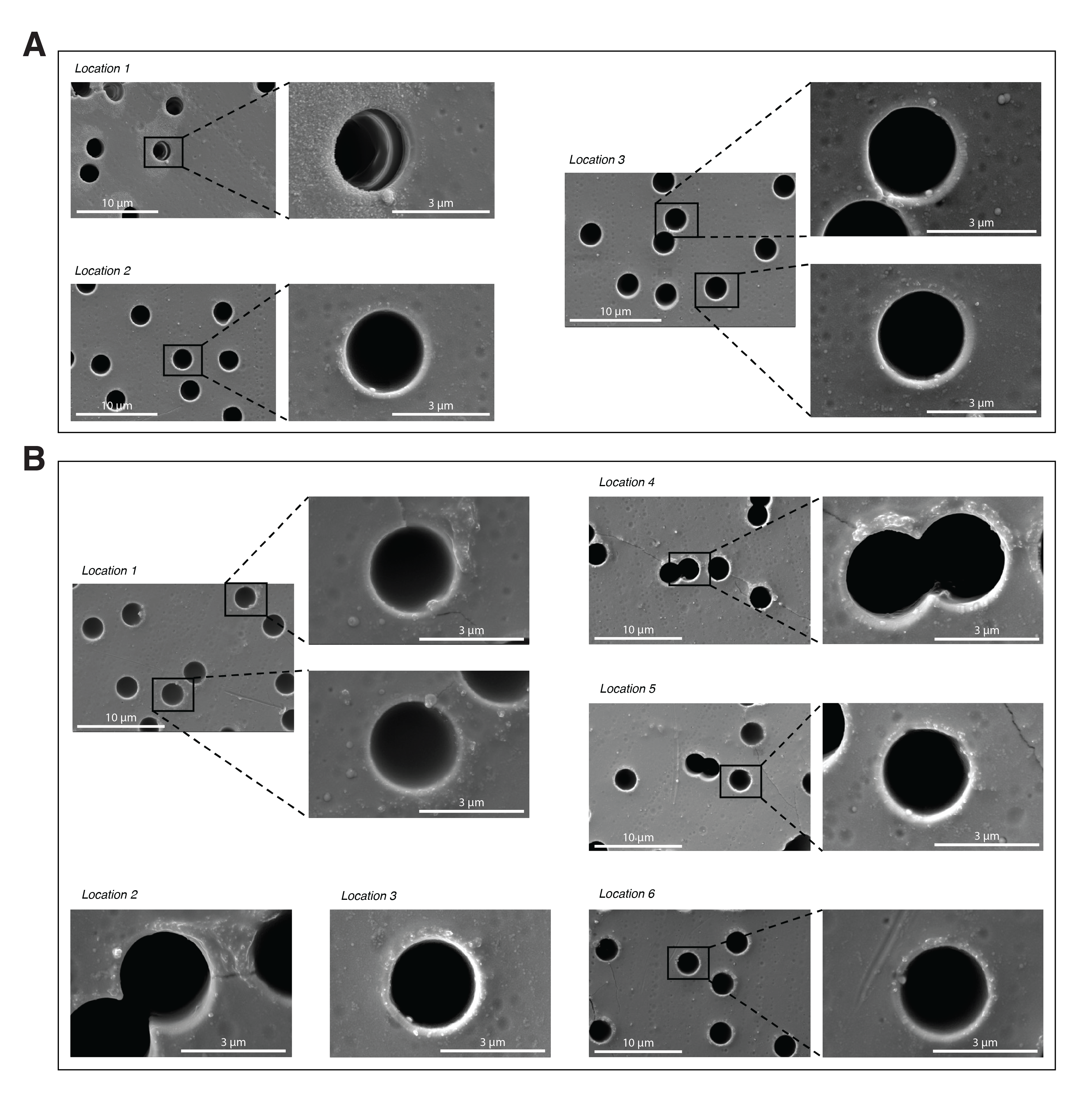

**Supplementary Fig. 4. Cryo-TEM micrographs of eluted EV-MNP complexes for the GluR2 (A) and GLAST (B) pulldowns demonstrating EVs bound to MNPs.** Abbreviations: cryo-TEM, cryo-transmission electron microscopy; EV, extracellular vesicle; MNP, magnetic nanoparticle.

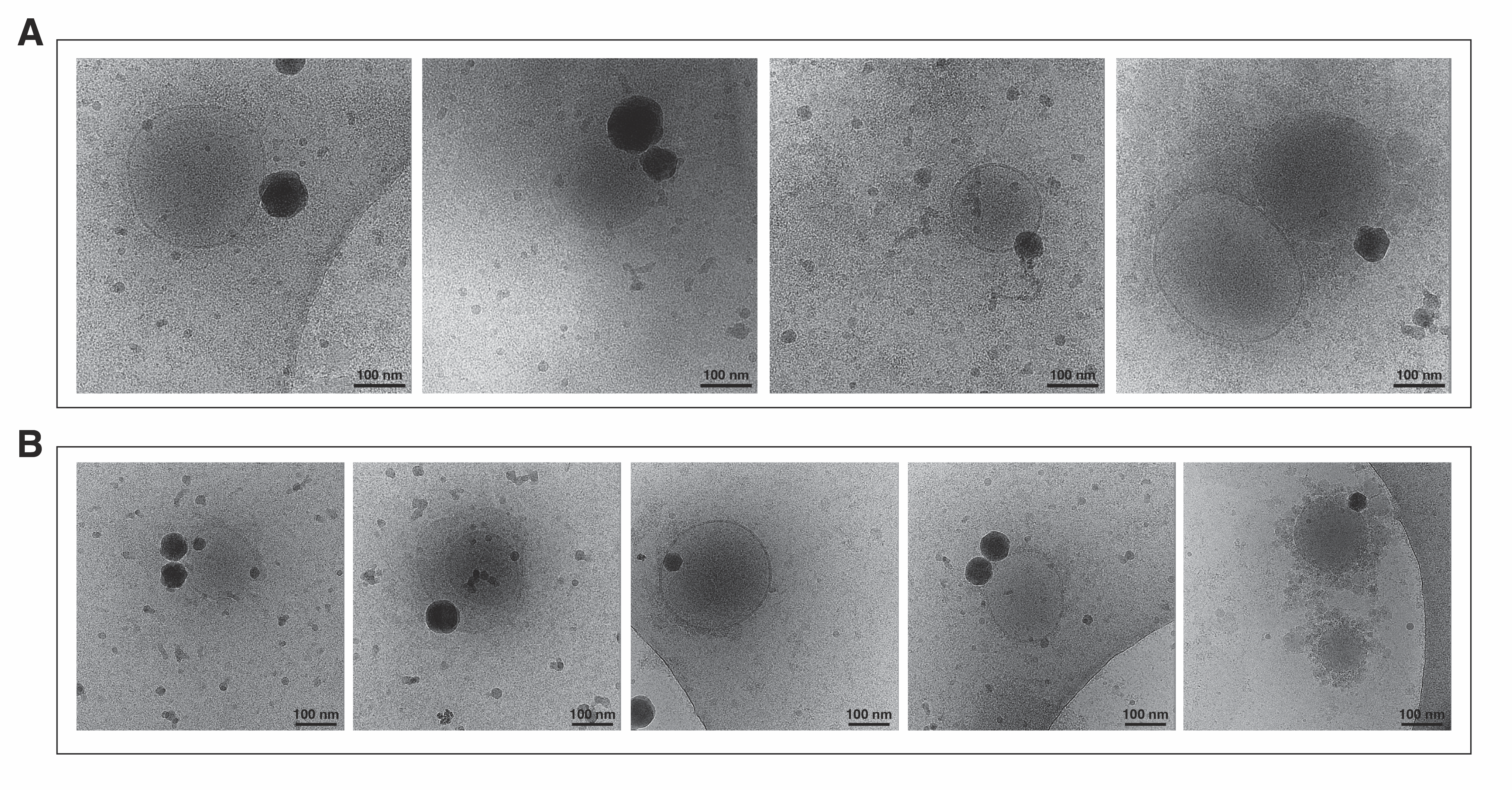

**Supplementary Fig. 5. SEM/HIM micrographs of eluted EV-MNP complexes for the GluR2 (A) and GLAST (B) pulldowns demonstrating EVs bound to MNPs.** Abbreviations: SEM, scanning electron microscopy; HIM, helium ion microscopy; EV, extracellular vesicle; MNP, magnetic nanoparticle.

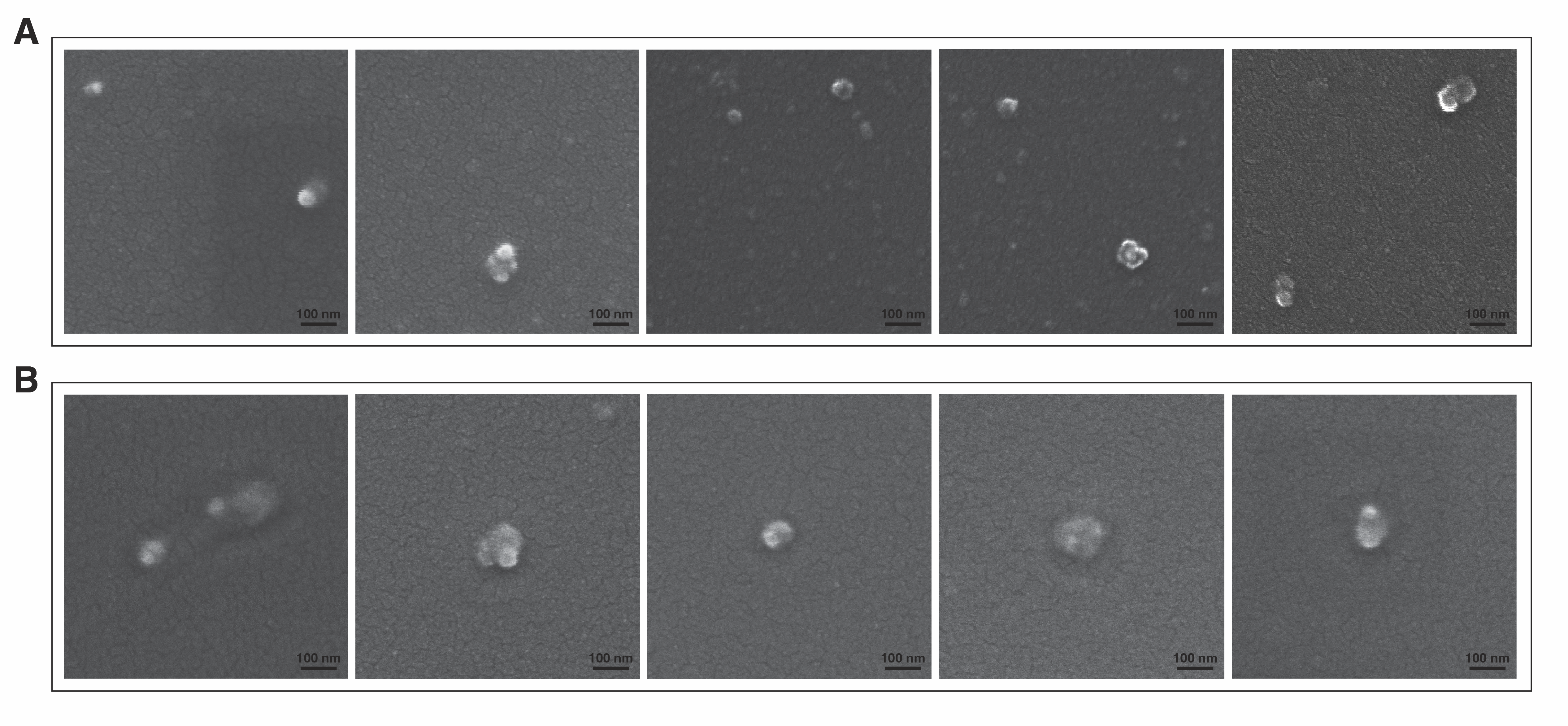

**Supplementary Fig. 6. NTA of EVs eluted off-chip from GluR2 (A) and GLAST (B) TENPO pulldowns from the pooled plasma of n = 8 patients per disease cohort.** Abbreviations: NTA, nanoparticle tracking analysis; EV, extracellular vesicle; TENPO, track-etch magnetic nanopore; LBD, Lewy body disease; AD, Alzheimer’s disease; AD/LBD, mixed Alzheimer’s and Lewy body disease; AD/ALB, Alzheimer’s disease with amygdala Lewy bodies.

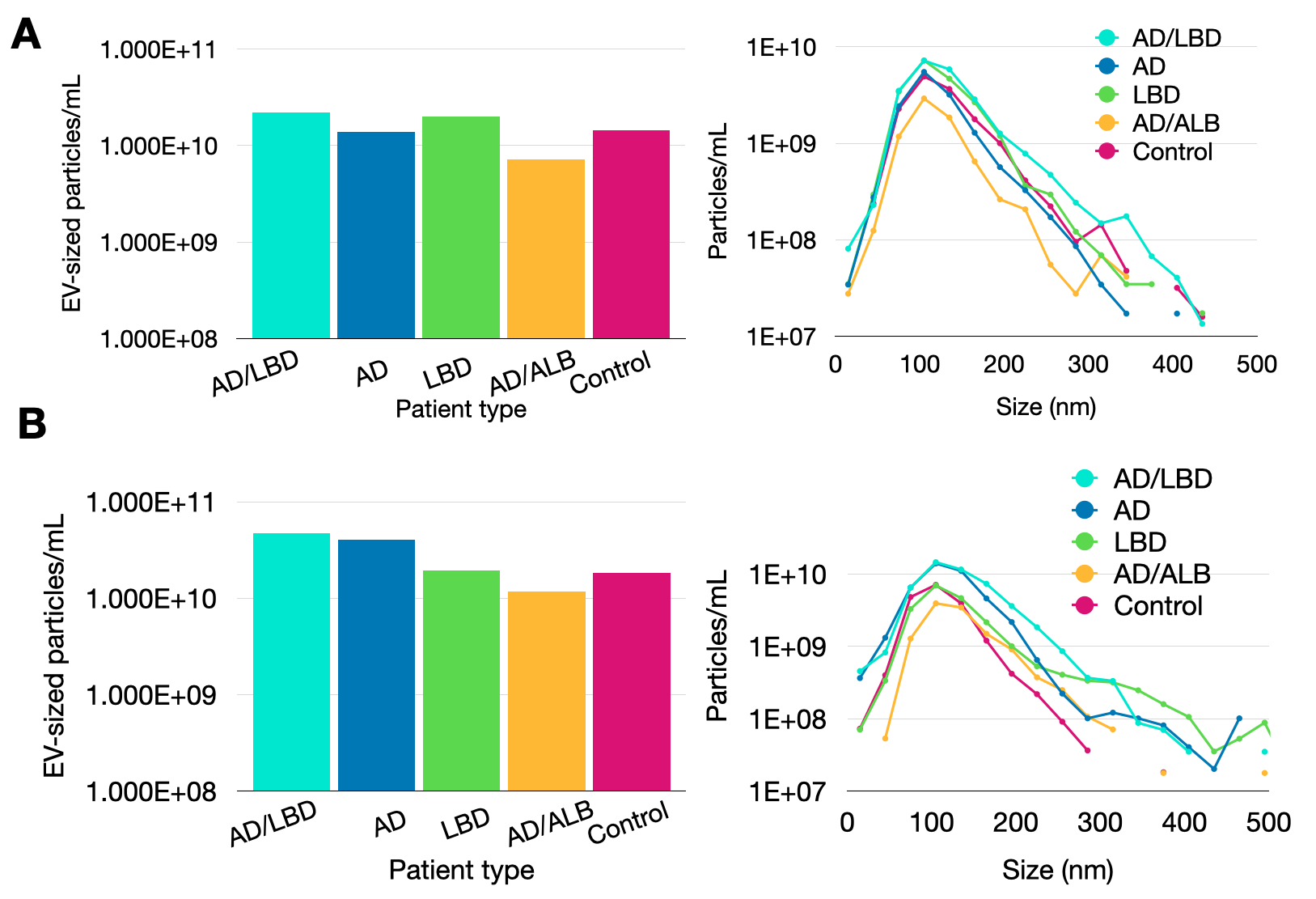

**Supplementary Fig. 7.** **Isotype control validation of GluR2 and GLAST antibodies**. Waterfall plots of ∆C_q_ values for a panel of miRNAs known to be expressed in CSF as a proxy for CNS-derived markers; miRNAs were measured in EVs isolated from plasma via TENPO devices using (A) GluR2-antibody-labeled MNPs versus isotype-labeled MNPs and (B) GLAST-antibody-labeled MNPs versus isotype-labeled MNPs. ∆C_q_ values were averaged across two replicates for GluR2 and three for GLAST. Markers are sorted by descending order of ∆C_q_ within the GluR2 compartment. Significance was assessed using a t-test; markers for which *P* value < .05 are highlighted in blue. Markers for which the C_q_ of both antibody and isotype were >37 were excluded; if a particular marker produced a ∆C_q_ within one compartment but was excluded from the other, it is shown with a red background. Abbreviations: CSF, cerebrospinal fluid; CNS, central nervous system; TENPO, track-etch magnetic nanopore; MNP, magnetic nanoparticle.

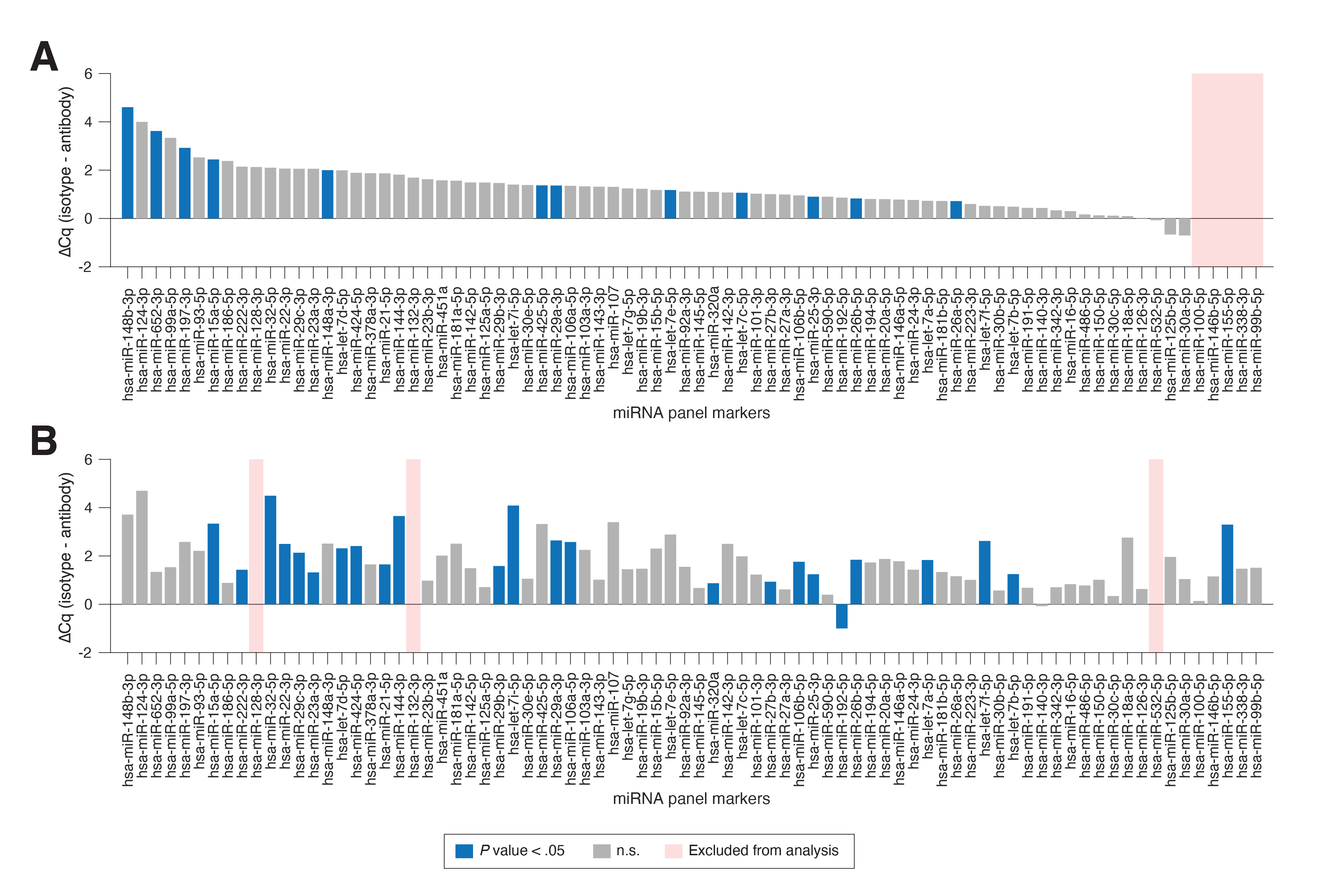

**Supplementary Fig. 8. NTA for GluR2 and GLAST TENPO pulldowns using mixed (combined neuron and astrocyte) (A) and astrocyte-only (B) cell culture media. (C) shows the NTA size distribution of anti-biotin MNPs alone.** Abbreviations: NTA, nanoparticle tracking analysis; TENPO, track-etch magnetic nanopore; MNP, magnetic nanoparticle.

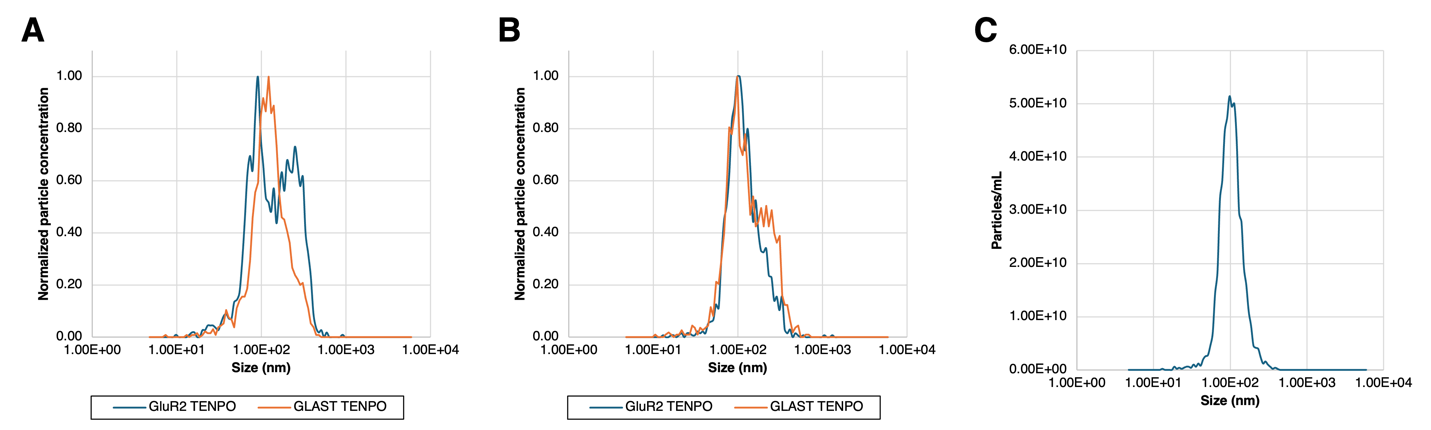

**Supplementary Fig. 9. Boxplots of relative expression of statistically significant GluR2 (A) and GLAST (B) nucleic acid and protein (C) biomarkers in comparing LBD versus AD.** Abbreviations: LBD, Lewy body disease; AD, Alzheimer’s disease; EV, extracellular vesicle; CPM, counts-per-million.

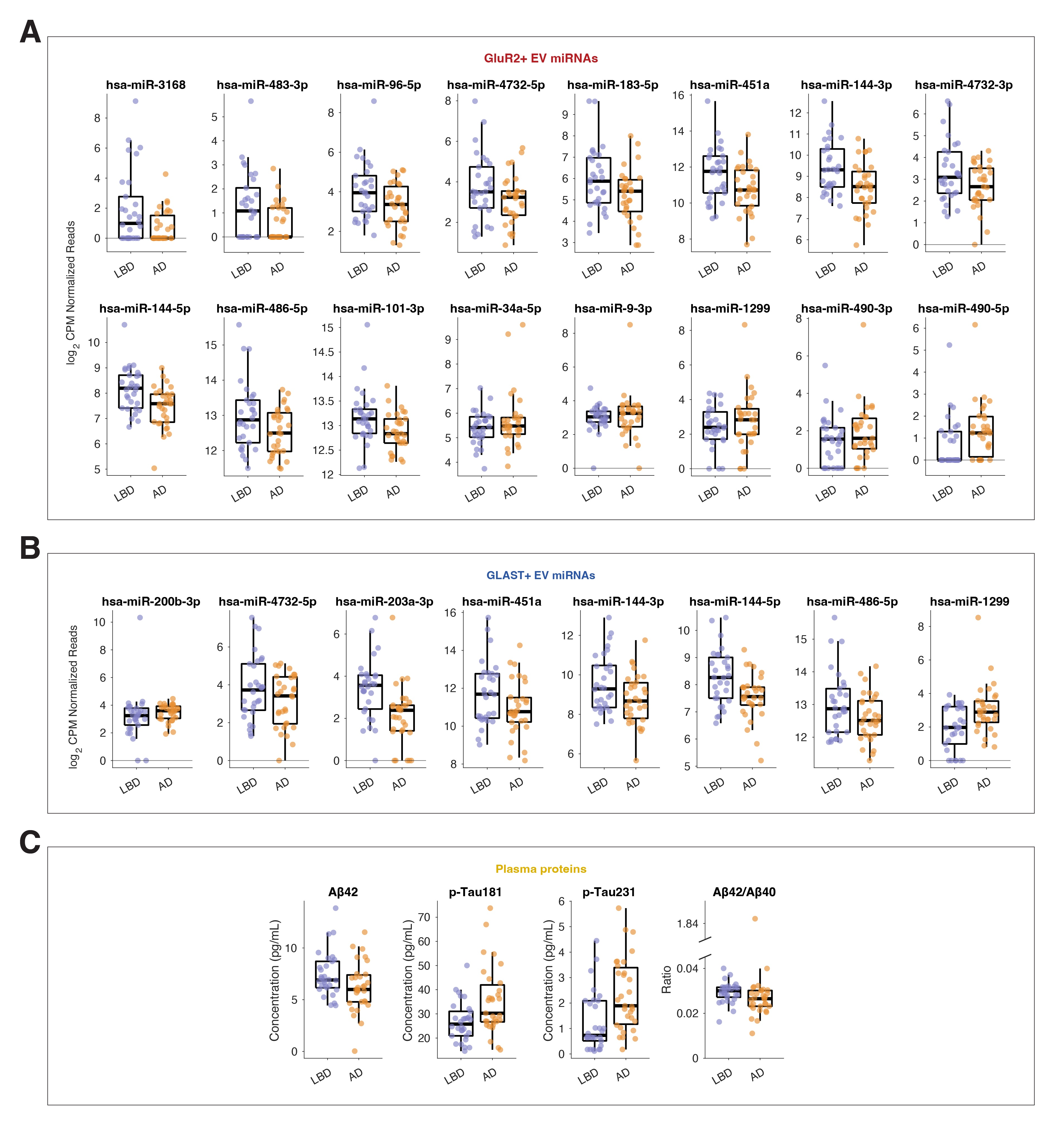

**Supplementary Fig. 10. ROC curves for Delong’s-significant differentially expressed GluR2+ EV miRNA (A), GLAST+ EV miRNA (B), and protein (C) biomarkers in comparing LBD versus AD. (D) shows correlation scatter plot for the top 30 markers based on AUC for the comparison of LBD versus AD.** Abbreviations: ROC, receiver operating characteristic; EV, extracellular vesicle; LBD, Lewy body disease; AD, Alzheimer’s disease; AUC, area under the curve.

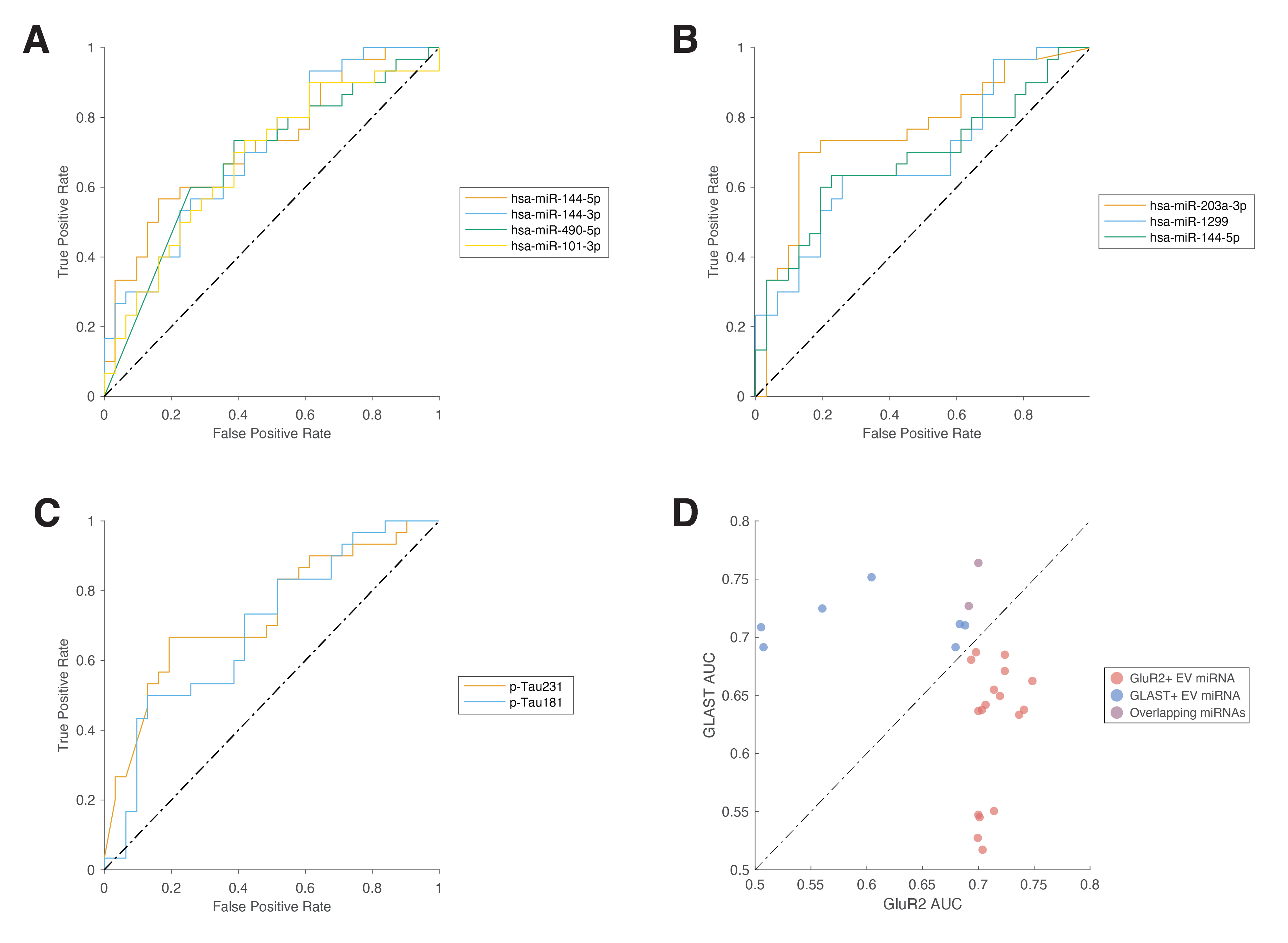

**Supplementary Fig. 11. Heatmaps of relative expression, volcano plots, and AUC graphs for the comparisons between dementia versus control.** (A) Heatmap of the z-score of log_2_(expression) for statistically significant biomarkers in a combined cohort of dementias (LBD, AD) versus control. (B) Volcano plot for biomarkers shown in panel A. (C) Top 30 biomarkers ranked by AUC for LBD + AD versus control. (D) Heatmap of the z-score of log_2_(expression) for statistically significant biomarkers in AD versus control. (E) Volcano plot for biomarkers shown in panel D. (F) Top 30 biomarkers ranked by AUC for AD versus control. (G) Heatmap of the z-score of log_2_(expression) for statistically significant biomarkers in LBD versus control. (H) Volcano plot for biomarkers shown in panel G. (I) Top 30 biomarkers ranked by AUC for LBD versus control. Error bars in C, F, and I represent standard error from bootstrapping 10x. The z-score color bar for A, D, and G, and the legend for the volcano plots (B, E, and H) and AUC graphs (C, F, I) are displayed on the right. Abbreviations: EV, extracellular vesicle; LBD, Lewy body disease; AD, Alzheimer’s disease; AUC, area under the curve; FDR, false discovery rate.

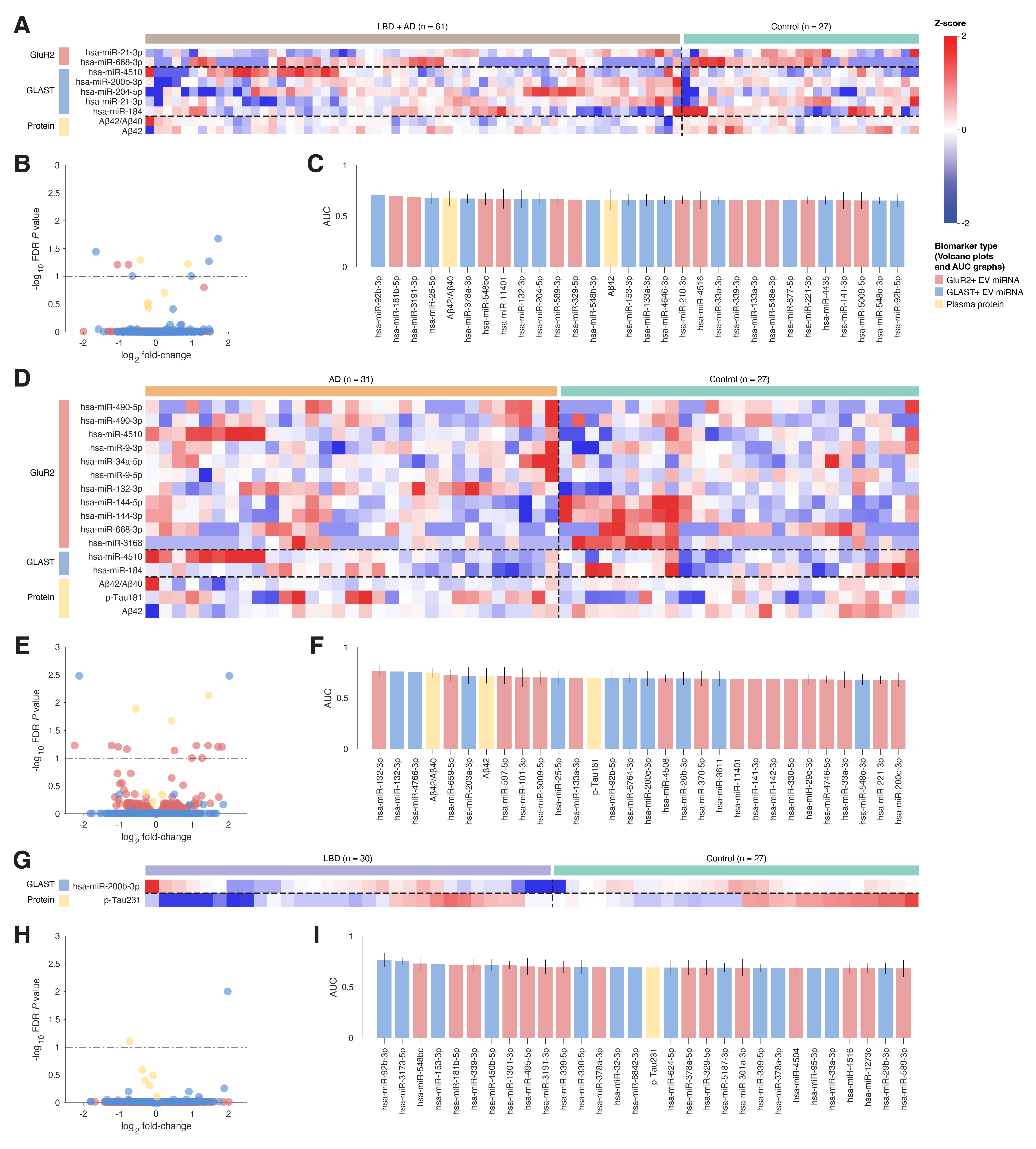

**Supplementary Fig. 12. Boxplots of relative expression for statistically significant biomarkers in comparing dementia versus control.** (A-C) Statistically significant biomarkers for the combined dementia (LBD + AD) versus control comparison. (D-F) Statistically significant biomarkers for the AD versus control comparison. (G-H) Statistically significant biomarkers for the LBD versus control comparison. Abbreviations: EV, extracellular vesicle; LBD, Lewy body disease; AD, Alzheimer’s disease; CPM, counts-per-million.

**Supplementary Fig. 13. ROC curves for the dementia versus control comparisons for Delong’s-significant differentially expressed biomarkers.** ROC curves for significant markers from the combined dementia (LBD + AD) versus control comparison for GLAST (A) and protein (B). ROC curves for significant markers from the AD versus control comparison for GluR2 (C) and protein (D). (E) ROC curve for the significant protein marker from the LBD versus control comparison. Abbreviations: ROC, receiver operating characteristic; LBD, Lewy body disease; AD, Alzheimer’s disease.

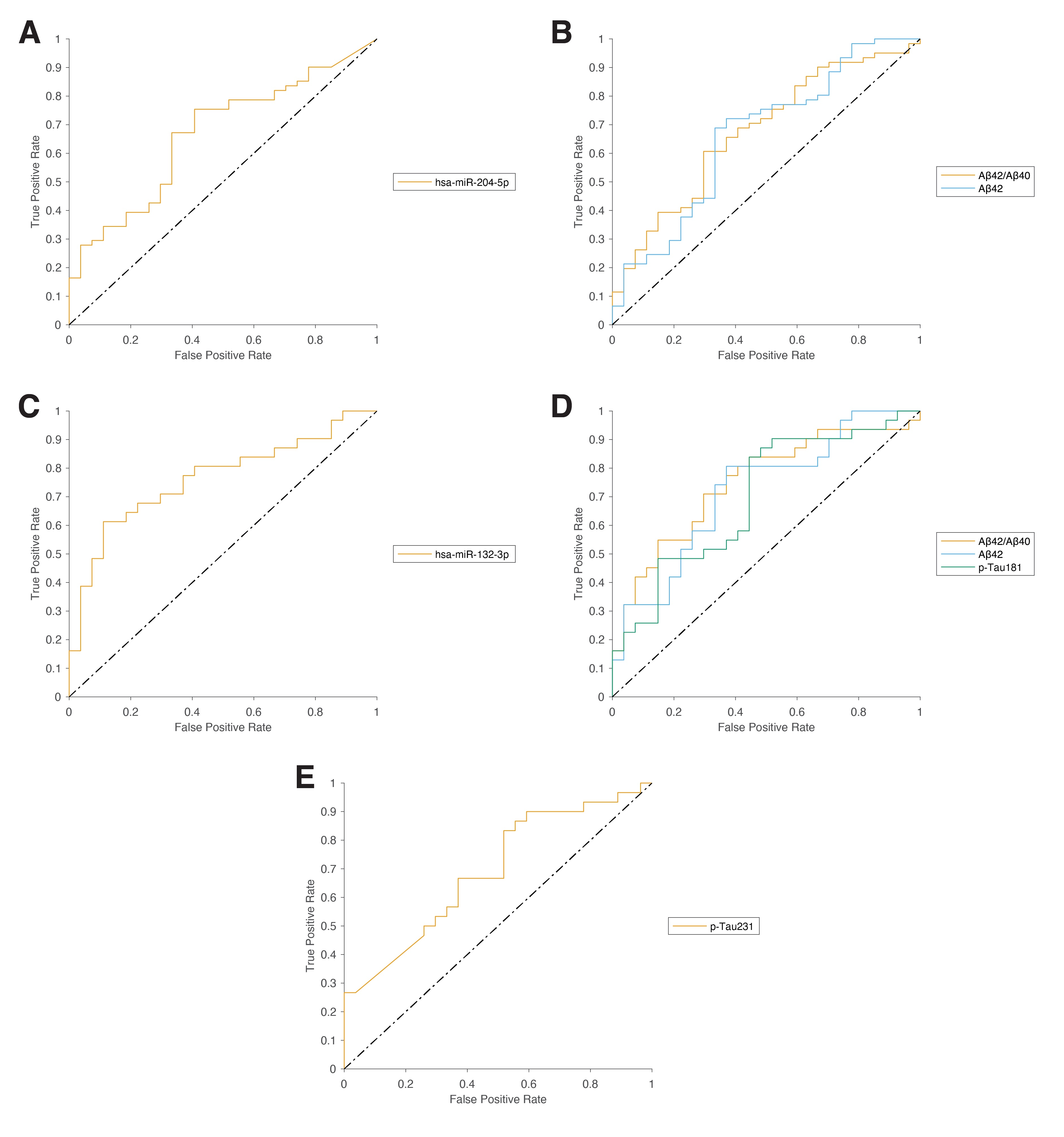

**Supplementary Table 4.** Significantly enriched (FDR *P* value < .05) GO BP terms associated with brain-related processes and neuroinflammation. Abbreviations: FDR, false discovery rate; GO, Gene Ontology; BP, biological process; TENPO, track-etch magnetic nanopore.

| TENPO Pulldown | GO BP Term | Target Genes | miRNAs | FDR *P* value |
| --- | --- | --- | --- | --- |
| GluR2 | nervous system development | 166 | 12 | 9.10038E-06 |
| GluR2 | brain development | 93 | 14 | 9.8826E-05 |
| GluR2 | neuron projection development | 54 | 13 | 0.00030806 |
| GluR2 | negative regulation of neuron apoptotic process | 49 | 13 | 0.01870682 |
| GluR2 | neuron migration | 43 | 13 | 0.00227707 |
| GluR2 | positive regulation of neuron projection development | 41 | 12 | 0.04425697 |
| GluR2 | negative regulation of neuron projection development | 35 | 12 | 6.5694E-06 |
| GluR2 | transport across blood-brain barrier | 34 | 10 | 0.00873633 |
| GluR2 | axonogenesis | 33 | 12 | 0.04123905 |
| GluR2 | neuron projection morphogenesis | 27 | 10 | 0.01894693 |
| GluR2 | response to cytokine | 25 | 12 | 0.00868455 |
| GluR2 | synapse assembly | 25 | 10 | 0.03586497 |
| GluR2 | positive regulation of neuron death | 19 | 9 | 0.04761377 |
| GluR2 | neuron apoptotic process | 18 | 11 | 0.01519507 |
| GluR2 | positive regulation of axonogenesis | 17 | 11 | 0.01623434 |
| GluR2 | positive regulation of axon extension | 17 | 11 | 0.02857206 |
| GluR2 | synaptic vesicle endocytosis | 17 | 10 | 0.02857206 |
| GluR2 | regulation of neuron projection development | 17 | 8 | 0.04675455 |
| GluR2 | maintenance of permeability of blood-brain barrier | 15 | 9 | 0.02446101 |
| GluR2 | neuronal stem cell population maintenance | 14 | 8 | 0.00565958 |
| GluR2 | axon extension | 14 | 11 | 0.01802768 |
| GluR2 | regulation of axon extension | 11 | 9 | 0.00145747 |
| GluR2 | synaptic vesicle transport | 10 | 8 | 0.02298226 |
| GluR2 | postsynaptic actin cytoskeleton organization | 8 | 7 | 0.00603828 |
| GluR2 | retrograde axonal transport | 8 | 6 | 0.01362023 |
| GluR2 | regulation of axon regeneration | 6 | 6 | 0.01894693 |
| GLAST | brain development | 45 | 7 | 0.00298682 |
| GLAST | negative regulation of neuron projection development | 17 | 5 | 0.00600164 |
| GLAST | response to cytokine | 14 | 6 | 0.02305511 |
| GLAST | neuron apoptotic process | 11 | 6 | 0.02078931 |
| GLAST | glial cell apoptotic process | 3 | 3 | 0.04040646 |

**Supplementary Table 5.** Significantly enriched (FDR *P* value < .05) KEGG pathways associated with the nervous system, neuron development, and brain-specific diseases. Abbreviations: FDR, false discovery rate; KEGG, Kyoto Encyclopedia of Genes and Genomes; TENPO, track-etch magnetic nanopore.

| TENPO Pulldown | KEGG Pathway | Target Genes | miRNAs | FDR *P* value |
| --- | --- | --- | --- | --- |
| GluR2 | Pathways of neurodegeneration - multiple diseases | 142 | 14 | 0.00319143 |
| GluR2 | Amyotrophic lateral sclerosis | 114 | 14 | 0.00118233 |
| GluR2 | Alzheimer disease | 111 | 14 | 0.01434632 |
| GluR2 | Parkinson disease | 74 | 13 | 0.0427041 |
| GluR2 | Axon guidance | 63 | 12 | 0.00012329 |
| GluR2 | Dopaminergic synapse | 53 | 12 | 3.9375E-05 |
| GluR2 | Neurotrophin signaling pathway | 48 | 13 | 2.9416E-05 |
| GluR2 | Spinocerebellar ataxia | 42 | 12 | 0.0344966 |
| GluR2 | Glioma | 33 | 12 | 0.00011778 |
| GluR2 | Long-term potentiation | 30 | 11 | 0.00019291 |
| GluR2 | Long-term depression | 24 | 9 | 0.00558459 |
| GLAST | Amyotrophic lateral sclerosis | 48 | 7 | 0.04728182 |
| GLAST | Axon guidance | 27 | 5 | 0.01768099 |
| GLAST | Neurotrophin signaling pathway | 23 | 6 | 0.0018131 |

**Supplementary Fig. 14. Interaction counts for top 10 individual genes targeted by differentially expressed GluR2+ EV (A) and GLAST+ EV (B) miRNA across the top 10 KEGG pathways in each pulldown.** Abbreviations: EV, extracellular vesicle; KEGG, Kyoto Encyclopedia of Genes and Genomes.

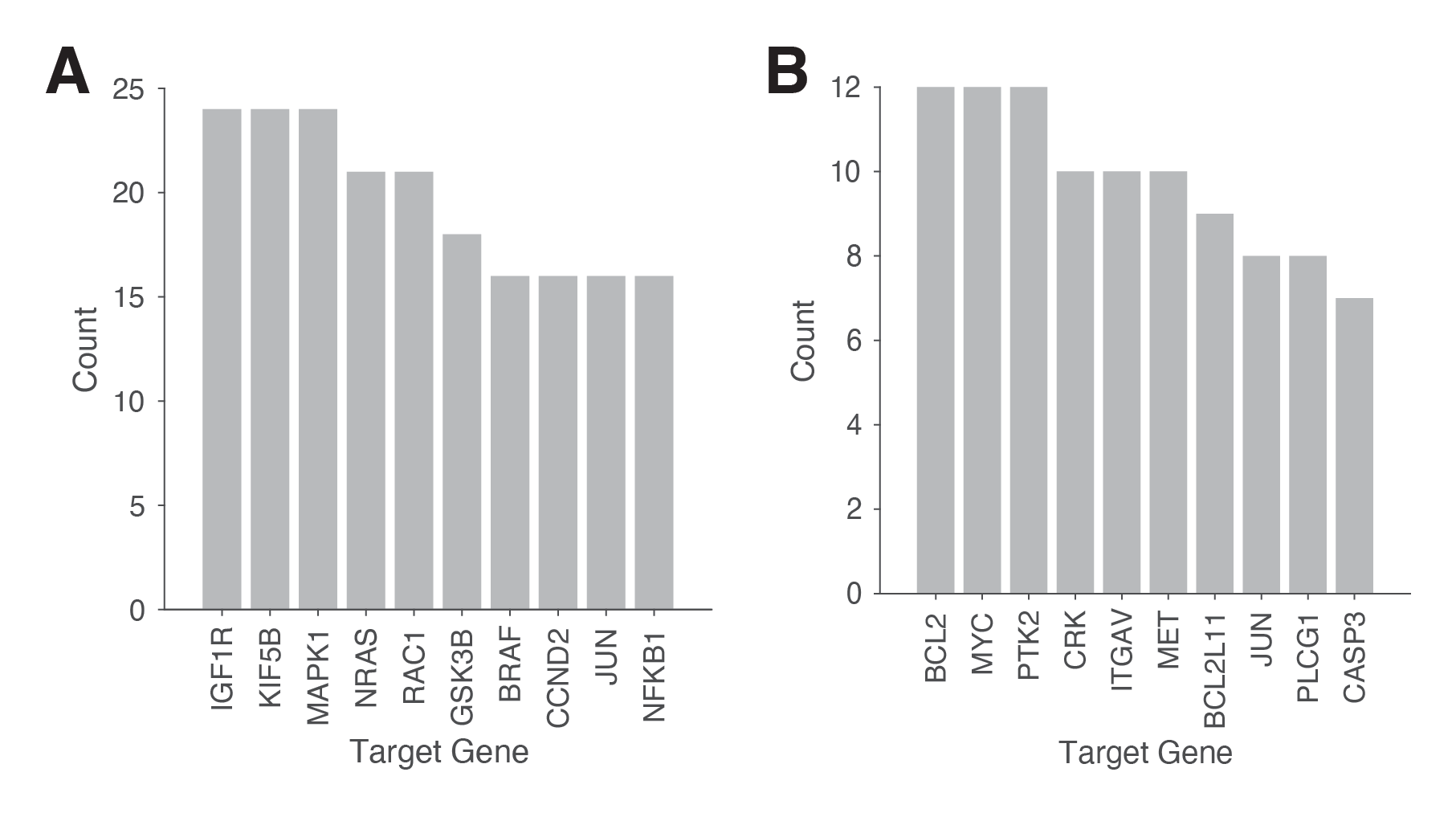

**Supplementary Table 6. Table of LASSO-selected markers with relevant literature references.** “N/A” indicates miRNAs for which literature references were not readily identifiable. Markers are listed in the order in which they were selected by LASSO. Abbreviations: LASSO, least absolute shrinkage and selection operator; AD, Alzheimer’s disease; CSF, cerebrospinal fluid; DLB, dementia with Lewy bodies; PD, Parkinson’s disease; Aβ, amyloid β; GFAP, glial fibrillary acidic protein; NfL, neurofilament light chain; FTD, frontotemporal dementia; ALS, amyotrophic lateral sclerosis; VaD, vascular dementia; PSP, progressive supranuclear palsy; CBD, corticobasal degeneration; PPA, primary progressive aphasia; MSA, multiple system atrophy; BBB, blood-brain barrier; MCI, mild cognitive impairment; EV, extracellular vesicle

| **Identified Marker** | **Related publications** |
| --- | --- |
| hsa-miR-548d-5p | - Plasma miR-548d is a potential biomarker for **schizophrenia**^1^ - miR-548d-5p overexpression is associated with poor prognosis in **glioma patients**^2^ |
| hsa-miR-26b-5p | - Serum hsa-miR-26b-5p discriminated **AD cases** with 89% sensitivity and specificity compared to controls^3^ - hsa-miR-26b is dysregulated in the blood, CSF, and brain tissue of **AD patients**^4-6^ |
| p-Tau231 | - p-Tau231 is a potential biomarker for assessing cognitive decline in **DLB** **patients**; DLB and PD patients had similar levels of plasma p-Tau231, which was lower than in AD patients and higher than in healthy controls^7^ - Plasma p-Tau231 showed the greatest measured change in patients with low Aβ burden within the **preclinical stage of AD**, as compared to other measured plasma biomarkers (p-tau181, p-tau217, GFAP, NfL and Aβ42/Aβ40)^8^ |
| hsa-miR-200c-3p | - miR-200c protected neurons from Aβ-induced damage in a **mouse model of AD**^9,10^ - miR-200c significantly decreases in abundance in **prion-infected** mice^11^ - Plasma miR-200c-3p is downregulated in C9orf72-associated **FTD and ALS patients compared to controls**^12^ |
| hsa-miR-330-5p | - miR-330-5p targets CDC42 which plays a role in hippocampal neuron dendritic spine formation; CDC42 is upregulated in the neurons of subjects with **AD versus controls**^13^ - miR-330-5p is implicated in regulation of **cerebral ischemia/reperfusion**^14^ |
| hsa-miR-6511a-3p | - N/A |
| hsa-miR-495-5p | - N/A |
| hsa-miR-132-3p | - miR-132 is known as “NeurimmiR” because it regulates many neurophysiological processes, including axon, dendritic and spinal maturation, and is commonly downregulated in **postmortem AD brain tissue**^15^ - Upregulation of miR-132-3p was found in **PD patients compared to controls**^16^ |
| hsa-miR-148b-5p | - Blood/serum hsa-miR-148b-5p is abnormally expressed in **AD patients compared to controls**^17,18^ |
| hsa-miR-339-5p | - miR-339-5p is reduced in brain tissue specimens of **AD subjects compared to controls**^19^ |
| hsa-miR-4748 | - N/A |
| hsa-miR-3611 | - N/A |
| hsa-miR-548as-3p | - N/A |
| hsa-miR-3173-5p | - Exosomal miR-3173-5p from plasma is significantly downregulated in **AD versus controls**^20^ |
| hsa-miR-5586-3p | - N/A |
| p-Tau181 | - p-Tau181 can distinguish **AD from FTD, VaD, PSP, CBD, PPA, PD, and MSA**^21^ |
| hsa-miR-424-5p | - miR-424-5p is a potential regulator of **BBB** permeability^22^ - hsa-miR-424-5p from plasma EVs is differentially expressed in **AD subjects compared to controls or subjects with VaD or MCI**^23^ |
| hsa-miR-6783-3p | - N/A |
| hsa-miR-548ap-5p | - N/A |
| hsa-miR-651-5p | - miR-651 is differentially expressed in the temporal cortex of **AD subjects**^24^ |

**Supplementary Fig. 15. ROC curves for the individual GluR2, GLAST, and protein biomarkers identified by LASSO for LBD versus AD, alongside the ROC curve for the 15-marker panel averaged across 5 repeats of 10-fold cross-validation.** Abbreviations: ROC, receiver operating characteristic**;** LASSO, least absolute shrinkage and selection operator; LBD, Lewy body disease; AD, Alzheimer’s disease.

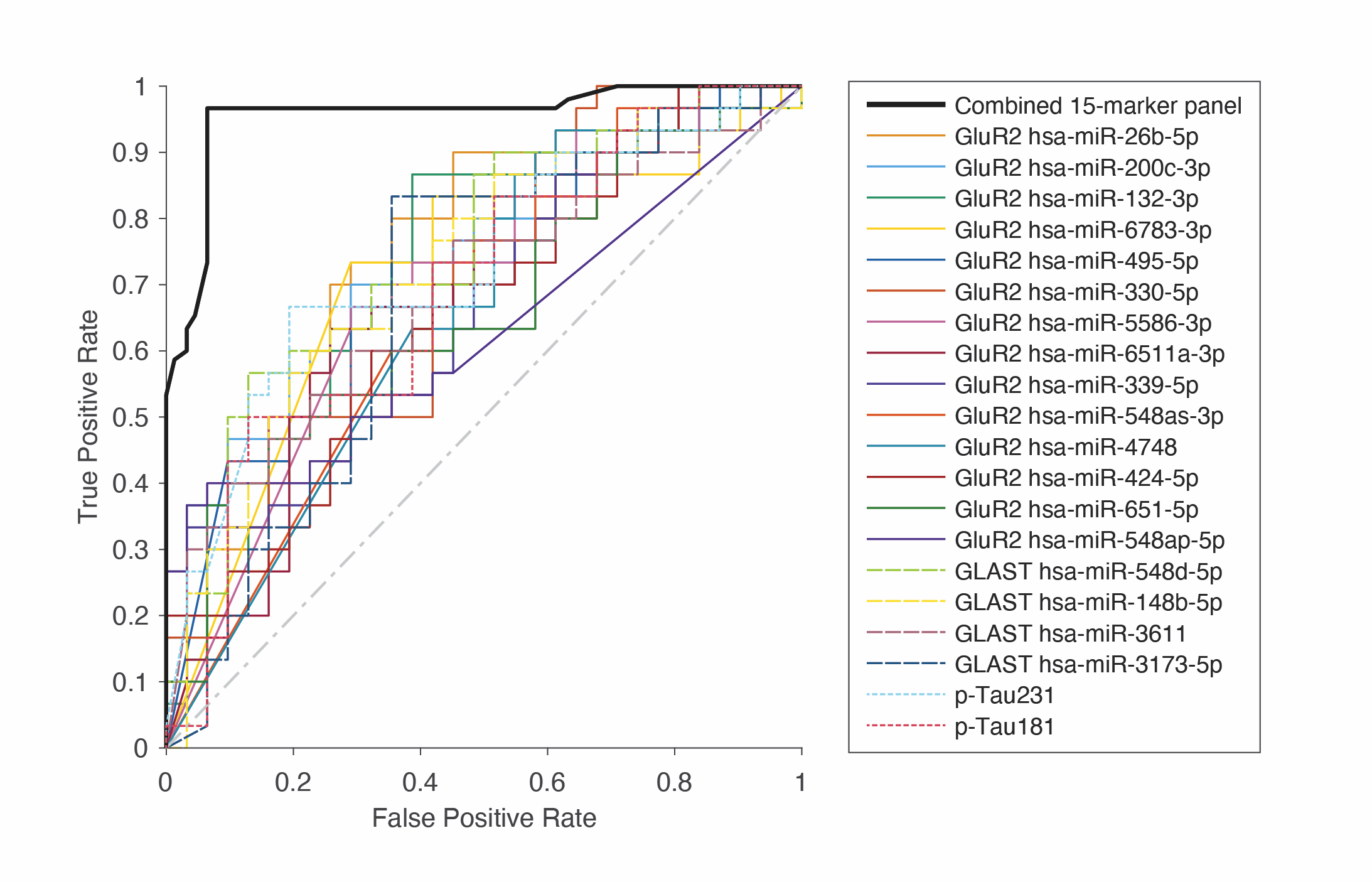

**Supplementary Fig. 16. Four-way boxplot comparisons between LBD, AD/LBD, AD/ALB, and AD for the markers identified via LASSO for LBD versus AD.** Abbreviations: LBD, Lewy body disease; AD, Alzheimer’s disease; AD/LBD, mixed Alzheimer’s and Lewy body disease; AD/ALB, Alzheimer’s disease with amygdala Lewy bodies; LASSO, least absolute shrinkage and selection operator; CPM, counts-per-million.

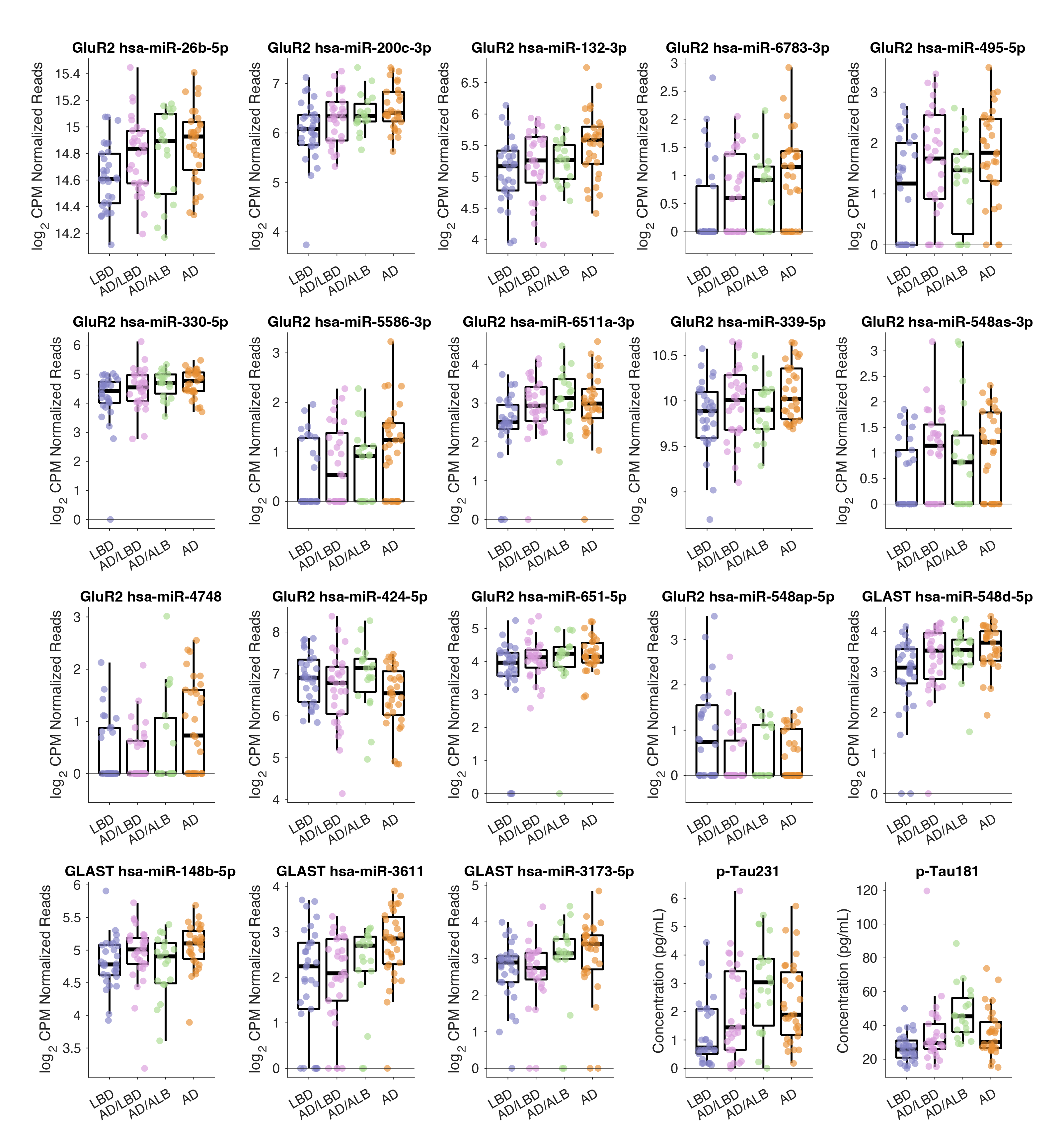
